# PHGDH is required for germinal center formation and is a therapeutic target in *MYC*-driven lymphoma

**DOI:** 10.1101/2021.11.11.468272

**Authors:** Annalisa D’Avola, Nathalie Legrave, Mylène Tajan, Probir Chakravarty, Ryan L. Shearer, Hamish W. King, Eric C. Cheung, Andrew J. Clear, Arief Gunawan, Lingling Zhang, Louisa K. James, James I. MacRae, John G. Gribben, Dinis P. Calado, Karen H. Vousden, John C. Riches

**Affiliations:** The Francis Crick Institute, 1 Midland Road, London, NW1 1AT, United Kingdom; Centre for Haemato-Oncology, Barts Cancer Institute, Queen Mary University of London, 3rd Floor John Vane Science Centre, Charterhouse Square, London EC1M 6BQ, United Kingdom; Centre for Immunobiology, Blizard Institute, Queen Mary University of London, London, UK; Metabolomics Science Technology Platform, The Francis Crick Institute, 1 Midland Road, London, NW1 1AT, U.K

## Abstract

The synthesis of serine from glucose is a key metabolic pathway supporting cellular proliferation in healthy and malignant cells. Despite this, the role that this aspect of metabolism plays in germinal center biology and pathology is not known. Here, we performed a comprehensive characterization of the role of the serine synthesis pathway in germinal center B cells and lymphomas derived from these cells. We demonstrate that upregulation of a functional serine synthesis pathway is a metabolic hallmark of B-cell activation and the germinal center reaction. Inhibition of phosphoglycerate dehydrogenase (PHGDH), the first and rate limiting enzyme in this pathway, leads to defective germinal formation and impaired high-affinity antibody production. In addition, overexpression of enzymes involved in serine synthesis is a characteristic of germinal center B-cell derived lymphomas, with high levels of expression being predictive of reduced overall survival in diffuse large B cell lymphoma. Inhibition of PHGDH induces apoptosis in lymphoma cells reducing disease progression. These findings establish PHGDH as a critical player in humoral immunity and a clinically relevant target in lymphoma.

## INTRODUCTION

B cells play a critical role in the humoral immune response that eliminates threats to the host by secreting highly specific antibodies. Activated B-cells can either differentiate into extrafollicular plasmablasts essential for early protective immune responses or enter a germinal center (GC). In the GC, B-cells undergo affinity maturation and eventually differentiate into plasma cells, which secrete high-affinity antibody critical to eliminate the infectious agent, or memory B cells that confer long-lasting protection from secondary infection.(1–4) In the GC reaction B cells are activated by antigen engagement via their B-cell receptor (BCR) and subsequent CD4^+^ T-cell help leading to MYC induction.(5) The cell-cycle regulator *MYC* is essential for the formation and maintenance of GCs.(6) *MYC* is commonly dysregulated in many high-grade B-cell malignancies including GC-derived lymphomas such as Burkitt lymphoma and diffuse large B-cell lymphoma (DLBCL).(7) *MYC* is a master regulator of metabolism, regulating the activity of many metabolic pathways including glycolysis and glutaminolysis. B-cell proliferation, either in the context of a GC-reaction or in B-cell lymphomas, requires significant alterations in cellular metabolism to sustain the demands of dividing cells.(8–10) B cells upregulate glycolysis following BCR-engagement, a metabolic switch that is also characteristic of many cancers, including high-grade lymphomas.(11–17) However, little is known about which metabolic pathways are involved in the utilization of glucose to support proliferating B cells.

One pathway that has emerged as a key metabolic node in cellular proliferation is the serine synthesis pathway (SSP). This uses a downstream product of glycolysis, 3-phosphoglycerate, to produce serine by the action of phosphoglycerate dehydrogenase (PHGDH), phosphoserine aminotransferase 1 (PSAT1) and phosphoserine phosphatase (PSPH).(18) Notably, all three of these enzymes are known MYC-targets.(19) Serine is necessary for glycine synthesis, phospholipid production, and acts as a one-carbon donor to the folate cycle, with serine-derived one-carbon units being used for the synthesis of purine nucleotides to support cell growth.(18, 20–22) Overexpression of SSP enzymes and increased serine biosynthesis from glucose is a feature of many types of cancer.(23) While some cancers acquire amplification or overexpression of *PHGDH*, the first and rate limiting step in this pathway, other types of cancers activate oncogenes such as *MYC, MDM2, KRAS* and *NRF2* leading to increased SSP enzyme expression.(19, 23–26) Upregulation of the SSP allows cells to increase *de novo* synthesis of serine from glucose when extracellular serine availability is limiting in the tumor microenvironment.(27–33) In support of this, a recent study has shown that inhibition of PHGDH is able to attenuate the growth of brain metastasis *in vivo*.(34–36) Despite this, little is known about the role of the SSP in B-cell lymphoma with no reports on its role in normal GC biology. Here, we performed a comprehensive characterization of the role of SSP in the GC reaction and lymphomagenesis. We reveal that upregulation of the SSP is a metabolic hallmark of B-cell activation and lymphoma, with PHGDH being a critical player in humoral immunity and a clinically relevant target in lymphoma.

## RESULTS

### Resting human naïve B cells lack expression of SSP enzymes which are induced upon activation

To understand the role of the SSP during B-cell responses *in vivo*, the expression of SSP genes in B-cell subsets isolated from reactive human tonsils from cancer-free individuals by singlecell RNA sequencing was analyzed.(37) We observed elevated expression of *PHGDH, PSAT1*, and *PSPH* in cycling GC B cells compared to naïve and non-proliferating B cells, suggesting an important role of the SSP in B-cell proliferation (Figure 1A). We then examined the expression of SSP enzyme proteins and mRNA in naïve B cells isolated from the peripheral blood of healthy individuals. While resting naïve B cells expressed very low-to-negligible amounts of PHGDH and PSAT protein, these enzymes were robustly induced 24-48 hours after stimulation by anti-IgM/G, CD40L and interleukin-4 (IL-4), signals that mimic those delivered *in vivo* to induce GC responses and B cell proliferation (Figure 1B-E). In contrast, PSPH was constitutively expressed, becoming further elevated after stimulation (Figure 1B-E). Treatment of naïve B cells by these stimuli alone or in combination revealed that upregulation of PHGDH and PSAT was predominantly driven by BCR-stimulation, which was synergistic with co-stimulation by CD40L and or IL-4 (Supplementary Figure 1A-B). The temporal dynamics of PHGDH and PSAT1 were noted to be different with PSAT1 expression being induced more rapidly than PHGDH, a pattern also observed in their mRNA transcripts (Figure 1E-F). Importantly, immunohistochemical (IHC) analysis of reactive human tonsils showed striking expression of PHGDH and PSAT1 within GCs, but not in marginal zone (MZ) areas (Figure 1G-H), confirming the upregulation of these enzymes as a hallmark of human GC B cells *in vivo*. We next assessed the dynamics of serine metabolism in activated B cells. We cultured isolated human B cells with U-[^13^C]-glucose and examined the steady-state incorporation of ^13^C-glucose-derived carbon into serine and glycine using LC-MS. While resting B cells fail to incorporate U-[^13^C]-glucose into serine, approximatively 50% of the intracellular serine pool was labeled from glucose in stimulated B cells, with 40% of serine carbon being fully labeled (Figure 1I). When serine is directly derived from fully labeled glucose it can be expected that all three of serine’s carbons will carry the ^13^C label (m+3). However, partially labeled serine isotopologues (m+1 and m+2) were also detected, likely due the inter-conversion of ^13^C-labeled and unlabeled serine and glycine (Figure 1I), indicating the bi-directional nature of this pathway. Taken together, these data indicate that resting human naïve B cells lack expression of SSP enzymes which are induced upon activation to provide a functional ability to synthesize serine and glycine from glucose.

**Figure 1.**
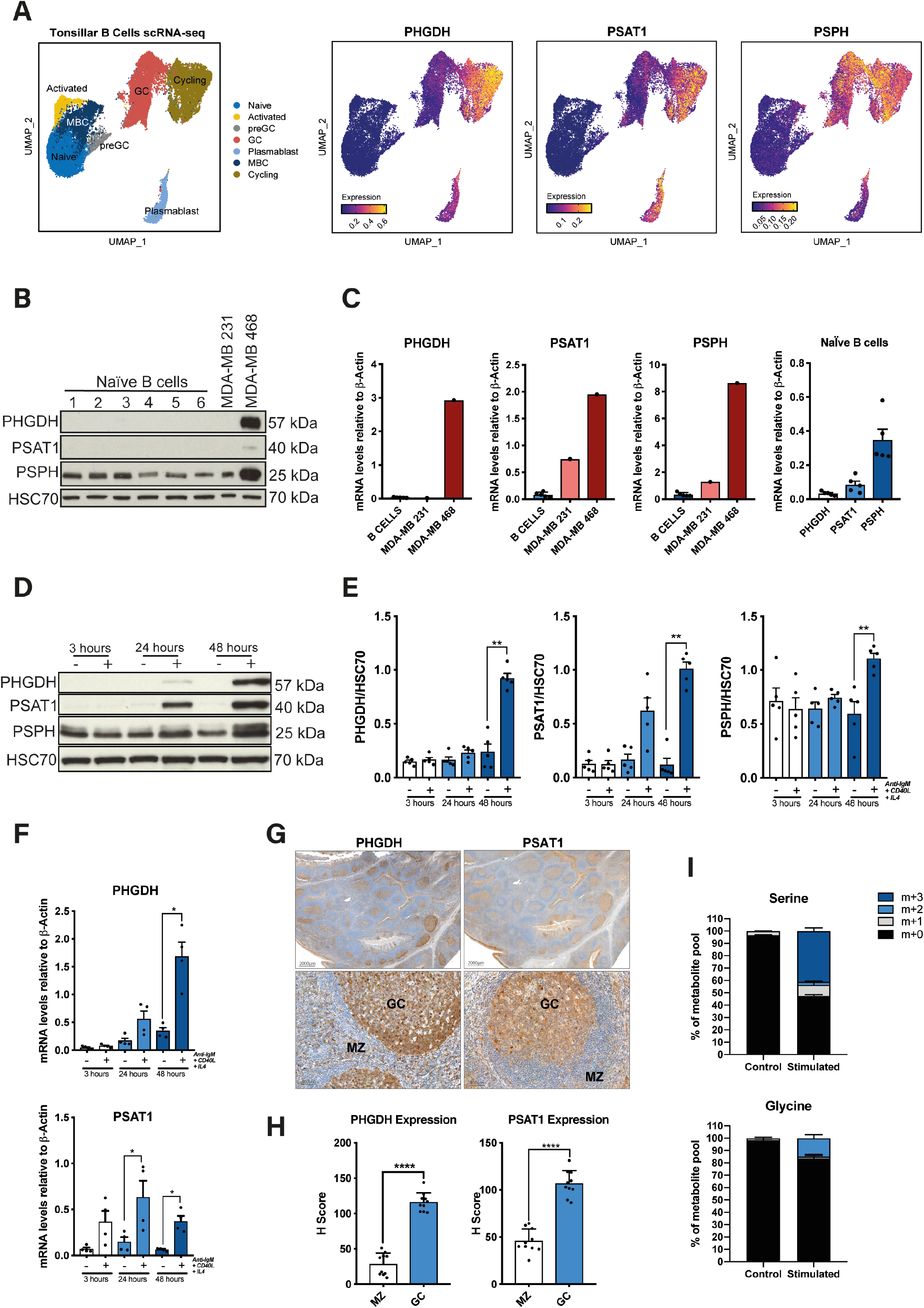
Upregulation of the SSP is a metabolic hallmark of GC B cells. (A) Uniform Manifold Approximation and Projection (UMAP) of tonsillar B cell scRNA clusters (including naïve, activated, pre-GC, total GC, plasmablasts, memory (MBC), and cycling B cells) (panel on the left). Expression of SSP-network genes in B-cell subsets (right). (B) Analysis of PHGDH, PSAT1 and PSPH protein levels in human naïve B cells isolated from blood bank volunteers by immunoblotting (n = 6). MDA-MB-231 and MDA-MB-468 cell lines were used as control for low- and high SSP-enzyme expression, respectively. (C) Quantification of specific transcript levels relative to b-actin mRNA levels. (D) Representative immunoblot of PHGDH, PSAT1 and PSPH proteins levels in resting and activated human naïve B cells. Human B cells were left unstimulated (-) or stimulated (+) with anti-IgM/G antibody, CD40 ligand (CD40L) and interleukin-4 (IL-4) for 3, 24 and 48 hours. (E) Quantification of protein levels shown in (D) normalized to HSC70. Individual samples (dots) and means (bars) values are plotted (n = 5; Mann Whitney test). (F) Relative mRNA expression of SSP enzyme genes in resting and activated human B cells determined by qPCR. Isolated human B cells were left unstimulated (-) or stimulated with (+) with anti-IgM/G antibody, CD40 ligand (CD40L) and interleukin-4 (IL-4) for 3, 24 and 48 hours before mRNA extraction. Specific transcript levels were determined relative to b-actin mRNA levels (n = 4; Mann Whitney test). (G) Representative immunohistochemical staining for PHGDH and PSAT1 in germinal center (GC) and marginal zone (MZ) areas in sections of human reactive tonsils and (H) quantification (n = 10; Mann Whitney test). (I) Mass isotopologue distribution of U-[^13^C_6_]-glucose-derived serine and glycine from human resting and activated B cells. B cells were left unstimulated (-) or stimulated (+) with anti-IgM/G antibody, CD40L and IL-4 for 48 hours. Cells were cultured for two hours in serine/glycine deplete media containing U-[^13^C_6_]-glucose.

### Characterization of the SSP in mice after activation *in vivo*

We characterized the expression of SSP enzymes in healthy resting murine B cells isolated from different tissue compartments. Consistent with the human data, there was low expression of PHGDH and PSAT1 protein in the peripheral blood, spleen and lymph nodes of wild-type mice (Figure 2A). IHC analysis of murine spleen and lymph nodes also confirmed low expression of SSP enzymes in resting B cells (Figure 2B-C). The expression of PHGDH and PSAT1 was assessed in murine spleen and lymph nodes by flow cytometry and IHC 8 days after immunization with sheep red blood cells (RBCs), a T-cell-dependent antigen that elicits robust GC responses. PHGDH and PSAT1 were specifically detected within peanut agglutinin^+^ (PNA^+^) GCs in the spleen (Figure 2B-C; Supplementary Figure 2A) and lymph nodes (Supplementary Figure 2B). Expression of SSP-involved enzymes was also increased in splenic B cells following *in vitro* stimulation for 24 hours (Figure 2D-F). Comparable to human B cells, activation of murine B cells resulted in a clear increase in their ability to synthesize serine and glycine from U-[^13^C]-glucose (Figure 2G).

**Figure 2.**
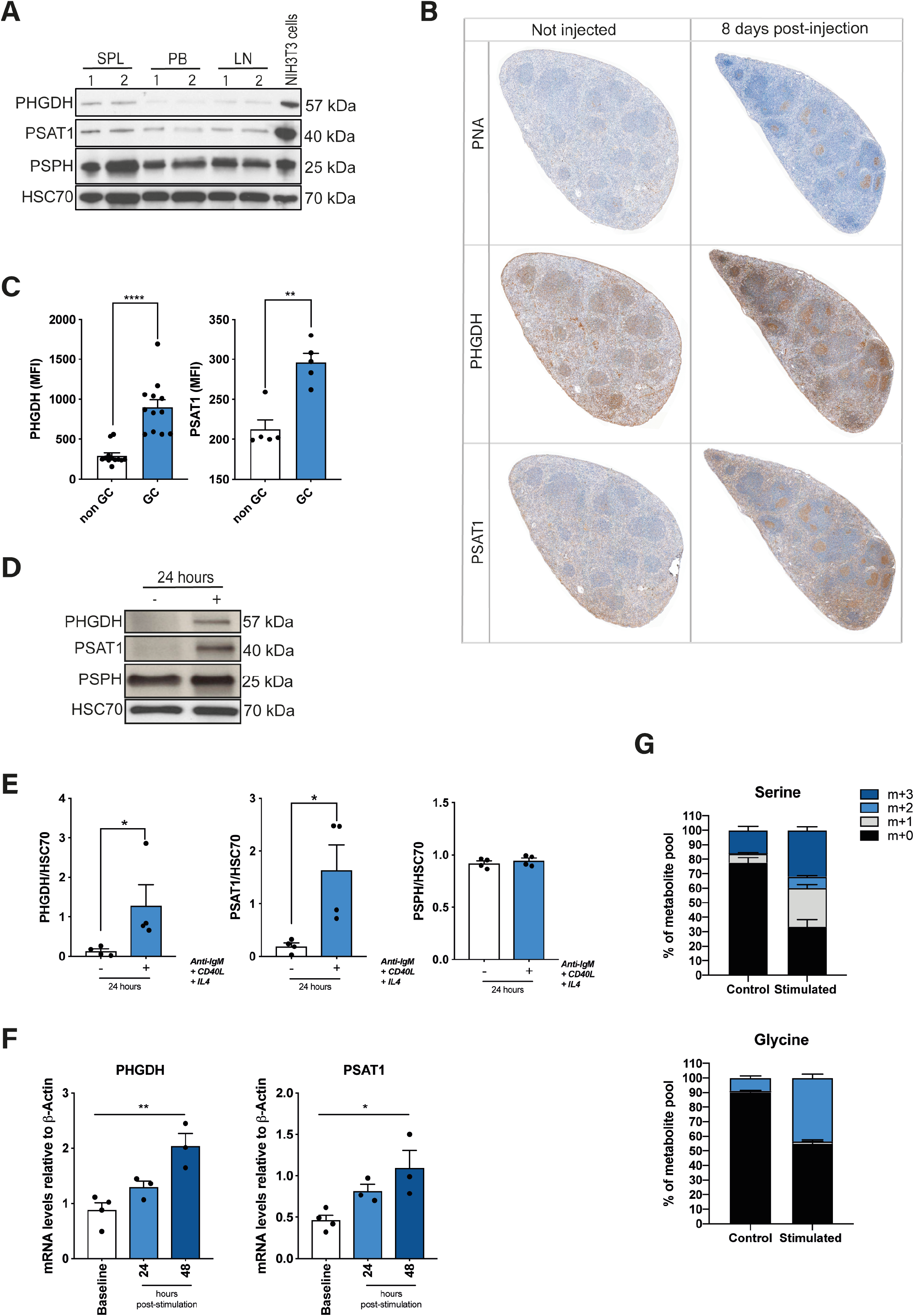
Characterization of the SSP in wild-type mice after activation *in vivo*. (A) Analysis of PHGDH, PSAT1 and PSPH protein levels in resting B cells isolated from mouse spleen (SPL), peripheral blood (PB) and lymph nodes (LN). NIH3T3 murine cells were used as control for high expression of SSP-related enzymes. (B) Representative immunohistochemical staining for PNA as GC marker, PHGDH and PSAT1 on consecutive spleen sections derived from mouse spleens 8 days after sheep RBC immunization. (C) Expression of PHGDH and PSAT1 in GC B cells and non-GC B cells harvested from mouse spleen 8 days after immunization with sheep RBC (Mann-Whitney test). (D) Representative immunoblots of PHGDH, PSAT1 and PSPH proteins levels in murine resting and activated B cells. Mouse B cells were isolated from spleen and left unstimulated (-) or stimulated (+) with anti-IgM/G antibody, CD40 ligand (CD40L) and interleukin-4 (IL-4) for 24 hours before protein extraction and (E) quantification of protein levels normalized to HSC70. Individual samples (dots) and means (bars) values are plotted (n = 4; Mann-Whitney test). (F) Relative mRNA expression of SSP enzyme genes in resting and activated mouse B cells as determined by qPCR. Isolated mouse B cells were left unstimulated (-) or stimulated with (+) with anti-IgM/G antibody, CD40 ligand (CD40L) and interleukin-4 (IL-4) for 24 and 48 hours before mRNA extraction. Specific transcript levels were determined relative to β-actin mRNA levels (n = 4; Mann-Whitney test). (G) Mass isotopologue distribution of U-[^13^C_6_]-glucose-derived serine and glycine from murine resting and activated murine B cells. Cells were left unstimulated (-) or stimulated (+) with anti-IgM/G antibody, CD40L and IL-4 for 48 hours. Cells were then cultured for two hours in media depleted from serine and glycine and containing U-[^13^C_6_]-glucose. ^13^C isotopologue distribution in serine and glycine was determined by LC-MS.

### Genetic loss and pharmacological inhibition of PHGDH impairs GC responses

To interrogate the role of the SSP in B-cell differentiation and GC responses, we targeted PHGDH, the first enzyme in the SSP, genetically and pharmacologically. We generated a conditional knockout murine model in which *Phgdh* was specifically deleted in B cells by crossing mice carrying floxed *Phgdh* alleles with mice expressing the Cre recombinase under control of the *Cd19* promoter (*Phgdh^fl/fl^;Cd19-Cre*). As expected B220^+^ B cells isolated from these mice did not show an increase in PHGDH expression following *in vitro* stimulation with anti-IgM/G antibody, CD40L and IL-4, whilst retaining the ability to upregulate PSAT1 (Supplementary Figure 3A). We then analyzed animals before and 8 days after immunization with sheep RBCs to assess the impact of deletion of *Phgdh* on GC responses (Figure 3A). Numbers of B220^+^CD38^+^Fas^+^ GC B cells were reduced in *Phgdh^fl/fl^;Cd19-Cre* mice following immunization, reflecting a reduction in both light zone (LZ) and dark zone (DZ) B cells (Figure 3B-C). These observations were also confirmed by IHC analyses showing a significant reduction in average PNA^+^ GC area and proportion of splenic sections occupied by PNA^+^ GCs in *Phgdh^fl/fl^;Cd19-Cre* mice (Figure 3D; Supplementary Figure 3B). Further confirmation of the B-cell specific nature of the *Phgdh* knockout was provided by a complete absence of PHGDH expression in GCs, in contrast to PSAT1, while PHGDH was expressed in the T-cell rich periarteriolar lymphoid sheaths (Supplementary Figure 3C). Thus, conditional knockout of *Phgdh* in B cells results in an impaired GC response.

**Figure 3.**
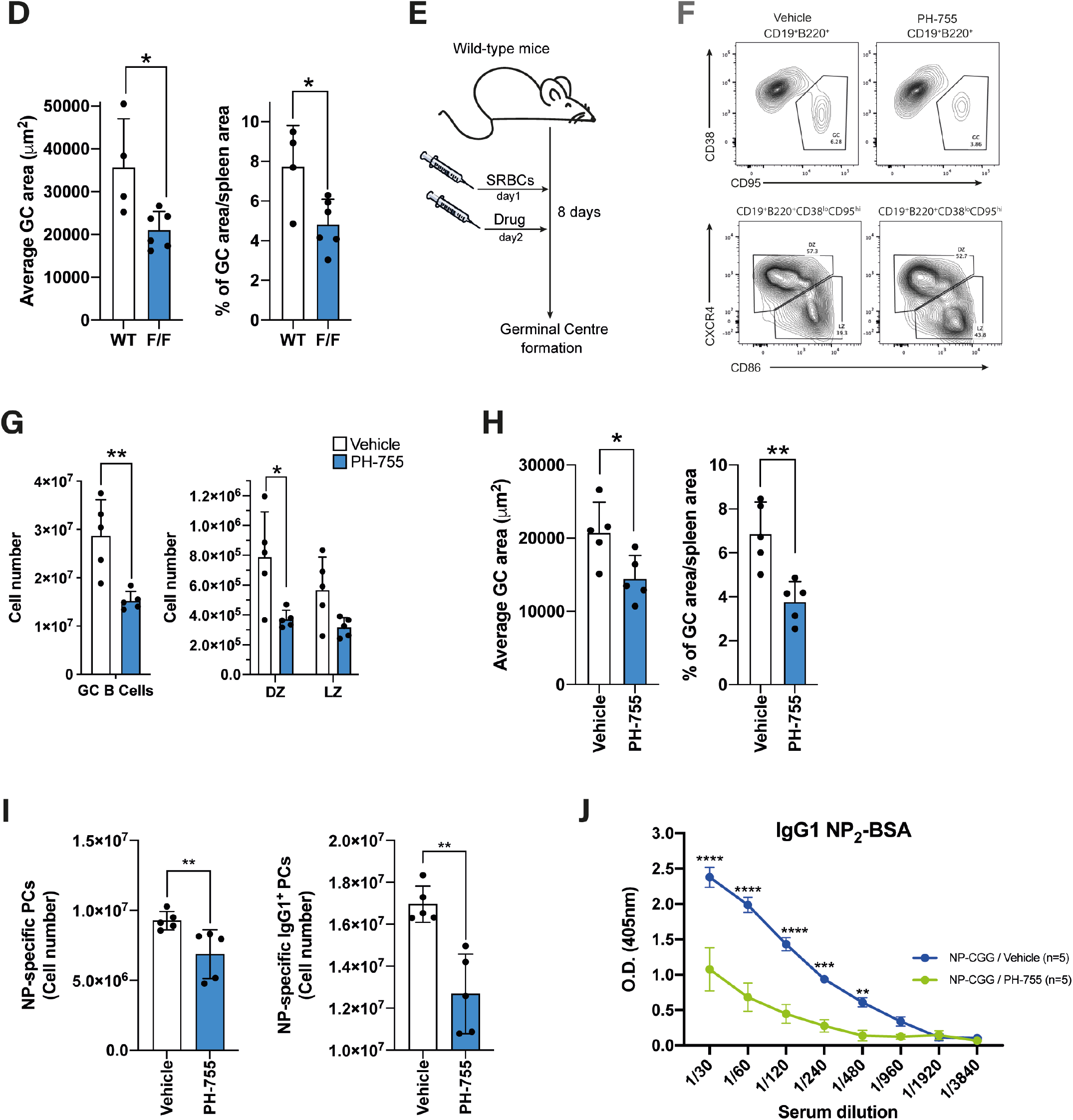
Genetic loss and pharmacological inhibition of PHGDH impairs GC responses. (A) Mice were injected with sheep RBCs and spleens examined by immunohistochemistry and flow cytometry 8 days after immunization with sheep RBCs. (B) Representative flow cytometric analysis of splenic B cells from either *Phgdh^+/+^;Cd19-Cre* (WT) or *Phgdh^fl/fl^;Cd19-Cre* (F/F) after immunization to identify GC B cells (CD19^+^B220^+^CD38^lo^CD95^hi^), and DZ (CD86^lo^CXCR4^hi^) and LZ (CD86^hi^CXCR4^l0^) B cells within GC splenic population. (C) Flow cytometric analysis of absolute number of B-cell subsets within CD19^+^B220^+^CD38^lo^CD95^hi^ splenic population from *Phgdh^fl/fl^;Cd19-Cre* (n = 6) and *Phgdh^+/+^;Cd19-Cre* (n = 4) after immunization. (D) Average GCs area (left) and proportion (%) of GC area per spleen area (right) from *Phgdh^fl/fl^;Cd19-Cre* (F/F) and *Phgdh^+/+^;Cd19-Cre* (WT) mice after immunization (Unpaired t-test)). (E) Wild-type mice were injected intra-venously with sheep RBCs one day before PH-755 treatment (300mg/kg PH-755 orally twice daily for 7 days). Spleens were collected for immunohistochemistry and flow cytometry 8 days after immunization. (F) Representative flow cytometric analysis of splenic B cells from mice injected with sheep RBCs for 8 days and treated with either vehicle or PH-755 to identify GC, DZ and LZ B cells within GC splenic population. (G) Flow cytometric analysis of absolute number of GC, DZ and LZ B cells within B220^+^ splenocytes collected from mice 8 days after sheep RBCs immunization; mice were treated with vehicle (n = 5) or PH-755 (n = 5)(Mann-Whitney test). (H) Average GCs area (left) and proportion (%) of GC area per spleen area (right) from vehicle- and PH-755-treated mice 8 days after sheep RBC immunization (Unpaired t-test). (I) Summary of NP_2_-specific plasma cells (PCs; left) and NP_2_-specific IgG1 PCs (right; total number per popliteal lymph nodes). Wild-type mice were treated with either Vehicle (n=5) or PH-755 (n=5) for 7 days. Animals were injected with NP-CGG one day before the beginning of PH-755 treatment. Popliteal lymph nodes were collected 8 days post NP-CGG immunization. (J) Serum antibody titers for NP_2_-specific IgG1 8 days post NP-CGG immunization.

We next assessed whether inhibition of GC responses could be replicated by pharmacological inhibition of PHGDH using a specific inhibitor, PH-755.(35, 36) Mice were injected with sheep RBCs 24 hours before starting treatment with PH-755 or vehicle control. GC responses were assessed by flow cytometry and IHC as before (Figure 3E). Treatment with PH-755 resulted in significantly reduced numbers of GC B cells comparable to that seen with the conditional knockout (Figure 3F-G). IHC analysis also showed a reduction in PNA^+^ GC area with an overall reduction in the proportion of splenic sections occupied by PNA^+^ GC B cells, but with preservation of PHGDH expression (Figure 3H, Supplementary Figure 3D-E). We then proceeded to assess the impact on NP-specific plasma-cell responses after NP-CGG immunization. PH-755 treatment resulted in a significant reduction of numbers of total NP-specific and NP-specific IgG1 plasma cells, which correlated with reduced titres of anti-NP IgG1 antibodies in the sera of these mice (Figure 3I-J) with no effect on NP-specific IgM responses (Supplementary Figure 3F). Taken together, these data show that PHGDH inhibition impairs GC-formation with a resultant reduction in high-affinity antibody production.

### PHGDH inhibition impairs B-cell proliferation and *de novo* serine and glycine synthesis

Based on previous observations showing the crucial role of the SSP in cell growth and proliferation, we proceeded to assess the impact of PHGDH inhibition on the behavior of primary stimulated murine B cells. Since GC B cells cannot survive *ex vivo* due to the rapid inception of a pro-apoptotic program,(38) we performed these experiments on B220^+^ cells isolated from spleens. Stimulated primary B cells from either *Phgdh^fl/fl^;Cd19-Cre* mice or *Phgdh^+/+^;Cd19-Cre* mice were tested for their ability to synthesize serine and glycine from glucose by incubating them with U-[^13^C]-glucose. Comparable experiments were performed with stimulated B cells taken from wild-type mice treated with either PH-755 or control. As expected, both genetic knockout and pharmacological inhibition of PHDGH almost completely abolished the ability of the cells to incorporate labeled glucose into serine at these time points (Figure 4A). We then assessed the impact of PHGDH inhibition on the proliferative capacity of B cells from *Phgdh^fl/fl^;Cd19-Cre* mice *in vitro* following stimulation, compared with *Phgdh^+/+^;Cd19-Cre* control B cells. B-cell proliferation measured by dye-dilution assay (Figure 4B) revealed only a slight reduction in proliferation of *Phgdh^fl/fl^;Cd19-Cre* B cells as compared to *Phgdh^+/+^;Cd19-Cre* cells when stimulated in serine-glycine replete medium (Figure 4C). However, in media lacking serine and glycine the proliferation of *Phgdh^fl/fl^;Cd19-Cre* B cells was completely abrogated whereas PHGDH induction following stimulation sustained the proliferation of *Phgdh^+/+^;Cd19-Cre* B cells (Figure 4B-C). Notably, the proliferative capacity of these PHGDH-deficient B cells could be partially rescued by the addition of glycine with formate as a one-carbon donor.(32) In addition, providing *Phgdh^+/+^;Cd19-Cre* B cells with formate and glycine allowed them to proliferate optimally in the absence of serine (Figure 4B-C). We then proceeded to investigate whether the changes in proliferation were accompanied by altered cell-cycling and/or apoptosis. Notably, the reduction in proliferation was mirrored by a decrease in the fraction of cells in S-phase, particularly when genetic knockout of *Phgdh* was accompanied by the removal of extracellular serine and glycine (Figure 4D). This decrease in cells in S-phase was accompanied by an accumulation of cells in G0/G1 phase (Figure 4D). Supplementing serine-starved *Phgdh^fl/fl^;Cd19-Cre* B cells with glycine and formate was able to partially rescue the fraction of cells in S-phase as reflected by cell proliferation (Figure 4D). We then proceeded to repeat these experiments with wild-type B cells cultured in various combinations of serine, glycine and formate, with PH-755 or vehicle control. Treatment with PH-755 largely recapitulated the pattern observed with the genetic ablation of *Phgdh*, with the exception that PH-755 treatment was able to partially inhibit B-cell proliferation even in the presence of extracellular serine and glycine (Figure 4E-G). Notably, we did not observe increased Caspase-3 activation as a marker of apoptosis in the *Phgdh* knockout or drug-treated B cells under any of the conditions (Figure 4H). In summary, these results show that inhibition of PHGDH effectively blocks *de novo* synthesis of serine and glycine from glucose which in turn has a cytostatic effect on primary murine B cells in the absence of extracellular serine and glycine.

**Figure 4.**
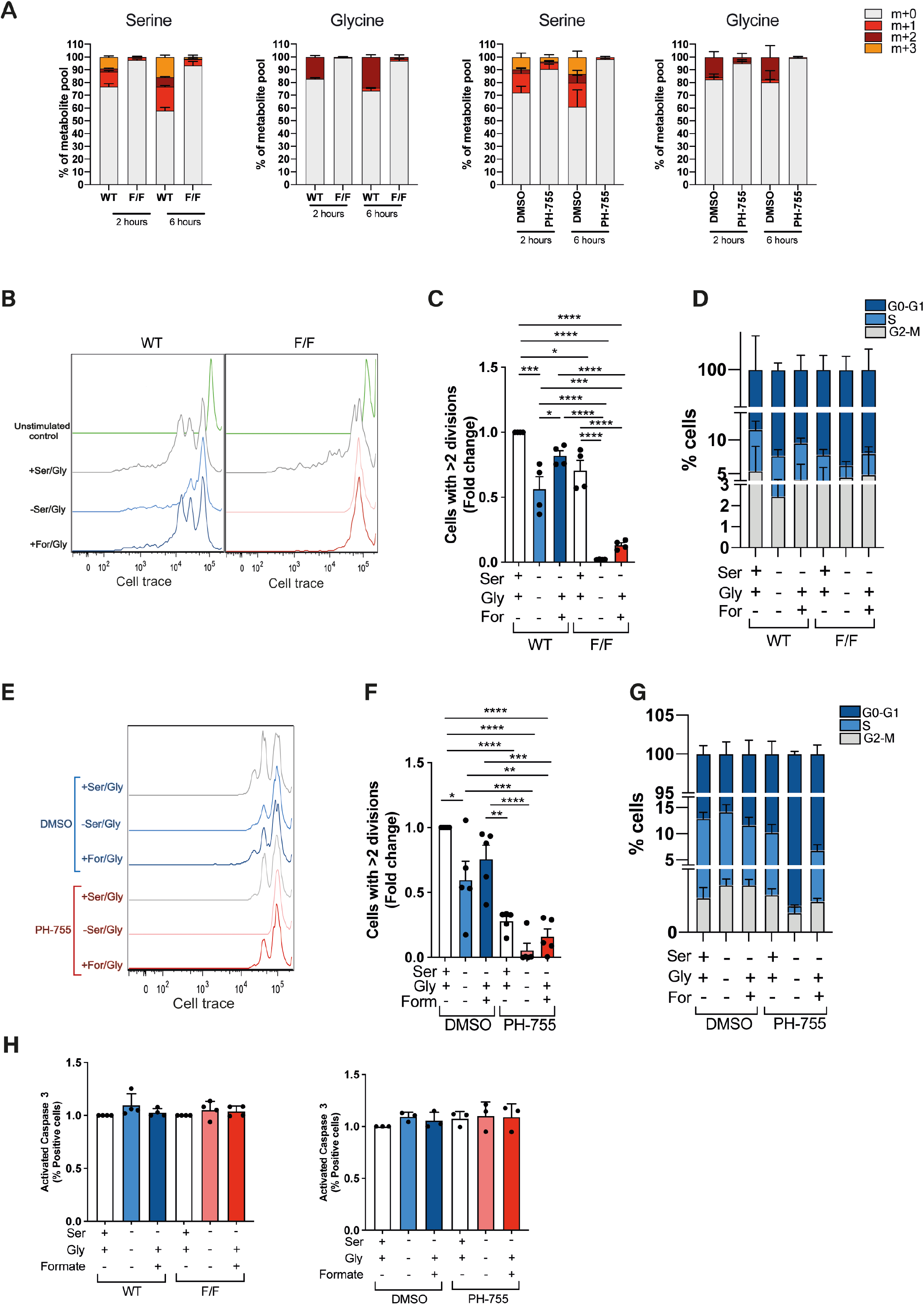
PHGDH inhibition impairs B-cell proliferation and *de novo* serine and glycine synthesis. (A) Mass isotopologue distribution of U-[^13^C_6_]-glucose-derived serine and glycine in B220^+^ B cells isolated from *Phgdh^+/+^;Cd19-Cre* (WT) or *Phgdh^fl/fl^;Cd19-Cre* (F/F) or wild-type BL6 mice. Isolated cells were stimulated with anti-IgM/G antibody, CD40L and IL-4 for 48 hours then cultured for two hours in serine/glycine-deplete media with U-[^13^C]-glucose (and treated with/out PH-755 for wild-type B cells). (B) Representative proliferation profiles and (C) quantification of B220^+^ B cells from either *Phgdh^+/+^;Cd19-Cre* (WT) or *Phgdh^fl/fl^;Cd19-Cre* (F/F) mice. Cells were labeled with the Cell Proliferation Dye eFluor 670, then cultured for 3 days with anti-IgM/G antibody, CD40L and IL-4 in complete media; serine/glycine-free medium; serine/glycine-free medium containing glycine and formate. (n=4 per genotype). (D) Cell-cycle analysis of B220^+^ cells isolated from *Phgdh^+/+^;Cd19-Cre* (WT) or *Phgdh^fl/fl^;Cd19-Cre* (F/F) mice. Cells were cultured for 48 hours with anti-IgM/G antibody, CD40L and IL-4 in complete media, serine/glycine-free medium or serine/glycine-free medium containing glycine and formate and then BrdU labeling and 7-AAD staining to assess cell-cycle. (E) Representative proliferation profiles and (F) quantification of B220^+^ B cells from wild-type mice. Cells were labeled with the Cell Proliferation Dye eFluor 670, then cultured for 3 days with anti-IgM/G antibody, CD40L and IL-4 in complete media, serine/glycine-free medium or serine/glycine-free medium containing glycine and formate in combination with DMSO or 10μM PH-755 (n=5 per group). (G) Cell-cycle analysis of B220^+^ B from wild-type mice. Cells were cultured for 48 hours with anti-IgM/G, CD40L and IL-4 in complete medium, serine/glycine-free medium or serine/glycine-free medium containing glycine and formate in combination with DMSO or 10μM PH-755 before cell-cycle analysis. (H) Activated caspase-3 on B220^+^ B cells from either *Phgdh^+/+^;Cd19-Cre* (WT) or *Phgdh^fl/fl^;Cd19-Cre* (left panel; n=4 per group) and treated as described in (D) and on B cells isolated from wild-type mice (right panel; n=3 per group) and treated as described in (G).

### Activation of SSP pathway in human MYC-driven GC lymphomas

One of the key emerging concepts in the cancer metabolism field over the last few years is that the metabolic phenotype of cancer cells reflects their cell-of-origin in combination with other factors such as oncogenic drivers and the tumor metabolic microenvironment.(39) In light of this we analyzed the role of the SSP in GC lymphomas. These were of particular interest due to the role of MYC in their pathogenesis and as a regulator of the SSP. The expression of PHGDH and PSAT1 was assessed by IHC in a series of diagnostic biopsies from patients with Burkitt lymphoma (BL), diffuse large B cell lymphoma (DLBCL) and chronic lymphocytic leukemia (CLL). Notably, very high expression of these two proteins was observed in BL consistent with a recent report,(40) with intermediate-to-high expression in DLBCL and relatively low expression in CLL (Figure 5A-B). Although biopsies from CLL patients showed the lowest expression, PHGDH and PSAT1 staining was significantly increased within proliferation centers, micro-anatomical sites in lymphoid tissues where CLL cells proliferate and where MYC is expressed (Figure 5C-D).(41, 42) Given the heterogeneity of expression in DLBCL patients, we next interrogated a published dataset (GSE10846) to investigate the relationship between SSP-gene expression and patient survival.(43) Importantly, high expression of *PSAT1* was significantly associated with poorer overall survival in DLBCL (Figure 5E) with a trend towards patients with high expression of *PHGDH* also having reduced overall survival. The expression of *PHGDH* and *PSAT1* in activated B-cell (ABC)-like or germinal center B-cell (GCB)-like DLBCL was also assessed due to the prognostic importance of these profiles.(44) There was no difference between the expression of *PHGDH* and *PSAT1* between these two subsets (Supplementary Figure 4A). This likely reflects our observations regarding the strong induction of these two enzymes upon activation of human B cells both *in vitro* and in GCs *in vivo* (Figure 1D-1H). Overall, the upregulation of the SSP is a feature of GC malignancies and can predict impaired overall survival in DLBCL.

**Figure 5.**
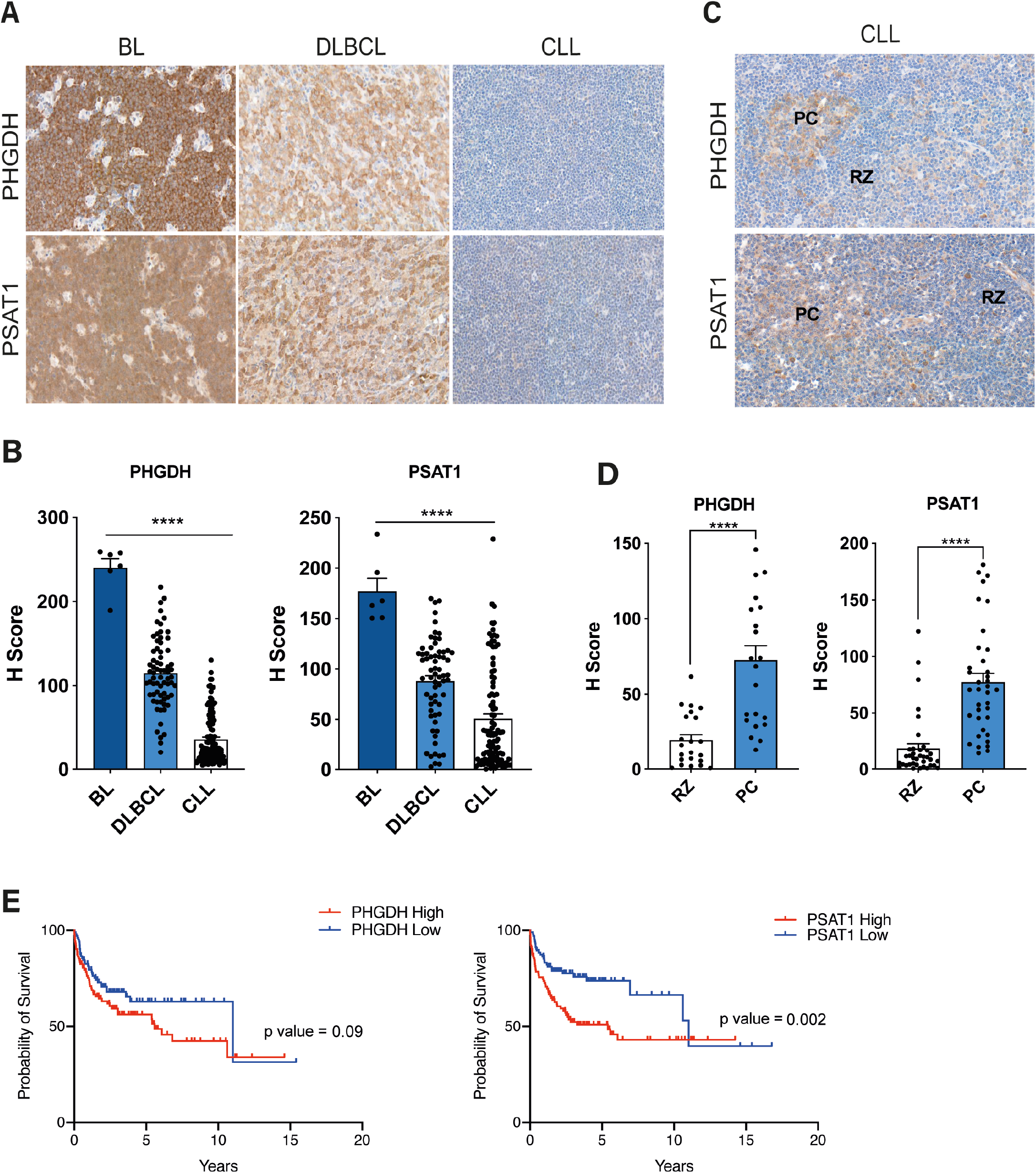
Human GC lymphomas are characterized by activation of the SSP pathway. (A) Representative immunohistochemical staining and (B) quantification for PHGDH and PSAT1 abundance in sections of human diagnostic biopsies from patients with Burkitt lymphoma (BL), Diffuse Large B-cell lymphoma (DLBCL) and Chronic lymphocytic leukemia (CLL). The statistical difference was analyzed using the ordinary one-way ANOVA. (C) Immunohistochemical analysis for PHGDH and PSAT1 in proliferation centers (PC) and resting zone (RZ) areas in sections from biopsies collected from patients with chronic lymphocytic leukemia (CLL) and (D) quantification. Individual samples (dots) and means (bars) values are plotted. The statistical difference was analyzed using the Mann-Whitney test. (E) Kaplan-Meier survival analysis of patients with DLBCL from a published dataset (GSE10846)(43). Patients whose *PHGDH/PSAT1* mRNA levels were within the top quartile were grouped as *PHGDH/PSAT1* high; those with *PHGDH/PSAT1* mRNA levels within the bottom quartile were grouped as *PHGDH/PSAT1* low. Statistical analyses were conducted with log-rank (Mantel-Cox) test.

### PHGDH inhibition impairs proliferation and promotes apoptosis in Burkitt lymphoma cells

We hypothesized that the SSP may represent a therapeutic target in human lymphoma. The expression of SSP enzymes was assessed in a panel of human lymphoma cell lines (Figure 6A; Supplementary Figure 5A). In contrast to other Burkitt lymphoma cell lines, Daudi cells had no expression of PHGDH, but did express PSAT1 and PSPH. Consequently, Daudi cells were unable to enter S-phase when cultured without extracellular serine and glycine, with cycling being partially rescued by glycine and formate. In contrast, Ramos and Raji cells were able to maintain the cell-cycle in the absence of serine and glycine (Figure 6B). However, when Ramos and Raji cells were treated with PH-755 to inhibit PHGDH they also became unable to cycle in the absence of serine and glycine, with PHGDH inhibition having no further effect on Daudi cells. In contrast to primary B cells, we observed significantly increased caspase-3 activation in Daudi cells cultured in serine/glycine-deplete conditions, and in all three cell lines when serine-glycine deprivation was combined with PHGDH inhibition with PH-755 (Figure 6C).

**Figure 6.**
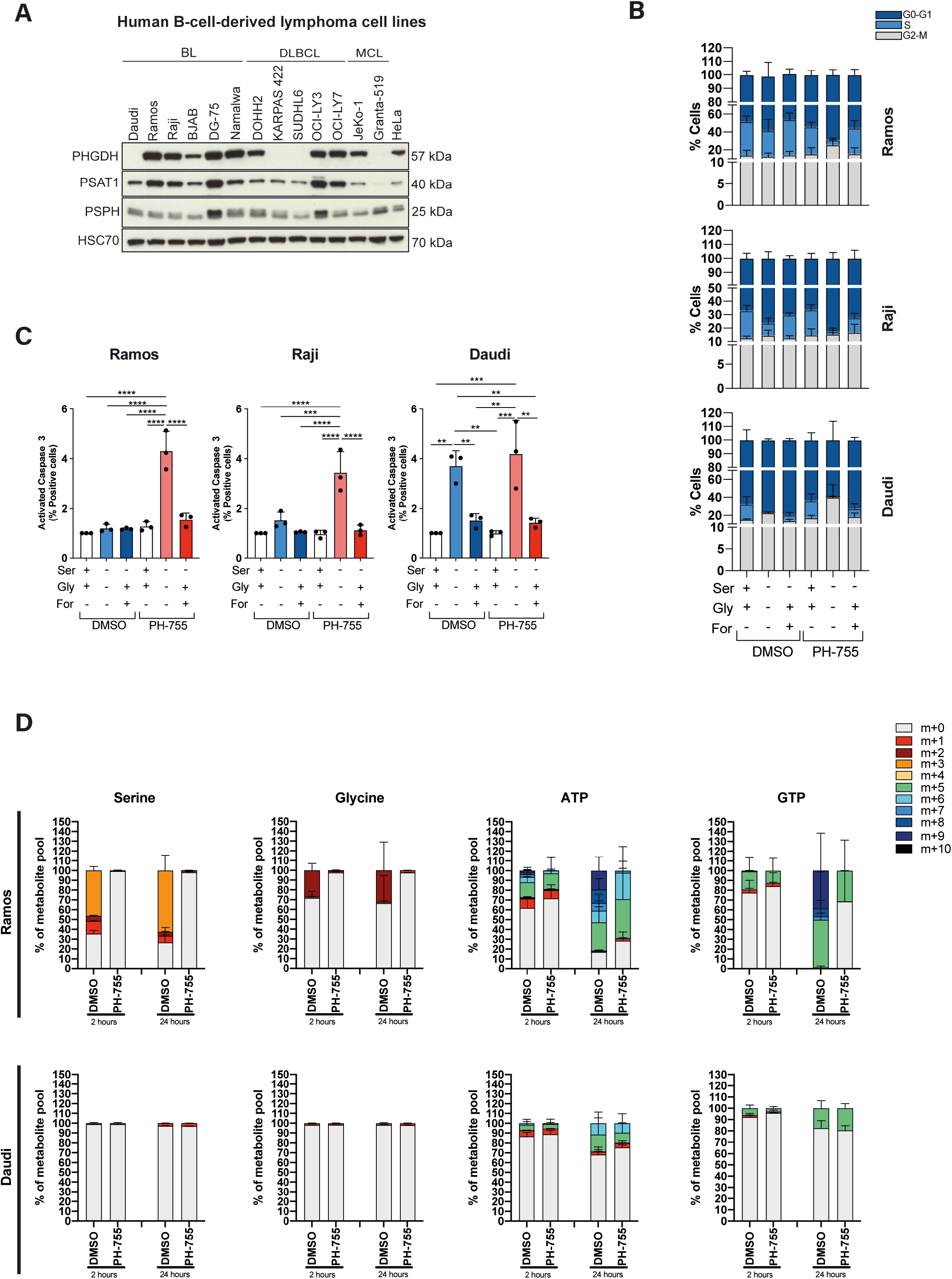
PHGDH inhibition impairs proliferation and promotes apoptosis in Burkitt lymphoma cells. (A) Western blot analysis of PHGDH, PSAT1 and PSPH protein expression in B cell-derived lymphoma cell lines (MCL=Mantle cell lymphoma). Representative of three independent experiments. HSC70 was used as loading control. (B) Cell cycle profile of Ramos (top), Raji (center) and Daudi (bottom) cells. Cells were plated either in complete medium or equivalent medium lacking serine and glycine supplemented or not with 0.5mM sodium formate and 0.4mM glycine and treated with DMSO (as a solvent control) or 10μM PH-755 followed incubation with 10μM BrdU and staining with anti-BrdU and 7-ADD. Data are presented as mean (± SEM) and are representative of three independent experiments, with value for DMSO-treated cells and cultured in complete medium set to 1.0. (one-way ANOVA with Tukey’s post hoc test). (C) Ramos (left), Raji (center) and Daudi (right) cells were cultured in the same conditions specified in (B) for 48 hours. Cells were then permeabilised, fixed, and stained for active Caspase-3. Positive cells for active Caspase-3 were analyzed by flow cytometry. Graph shows the mean (± SEM) derived from three independent experiments, with value for DMSO-treated cells and cultured in complete medium set to 1.0. (one-way ANOVA with Tukey’s post hoc test). (D) Mass isotopologue distribution of U-[^13^C_6_]-glucose-derived serine and glycine for Ramos (top) and Daudi (bottom) cells cultured for 2 and 24 hours in medium lacking serine and glycine in presence of U-[^13^C_6_]-glucose (10mM) and treated with DMSO or 10μM PH-755. Serine, Glycine, ATP and GTP levels were measured by LC-MS. The percentage distribution of each isotopologue for their respective metabolite pool is shown. Data are presented as mean (± SEM) of six repeats and are representative of three independent experiments.

We assessed the ability of these cell lines to synthesize serine, glycine and the purine nucleotides adenosine triphosphate (ATP) and guanine triphosphate (GTP) from glucose. The cell lines were cultured with U-[^13^C]-glucose for 2-24 hours in the absence of serine and glycine with labeling being assessed by LC-MS. Ramos and Raji cells grown in the absence of serine and glycine diverted glucose into *de-novo* serine and glycine synthesis which was prevented when treated with PH-755 (Figure 6D; Supplementary Figure 5B). Furthermore, labeling of higher isotopologues (≥m+6) of ATP and GTP demonstrated that these nucleotides were being synthesized via the SSP, which could again be inhibited by PH-755 (Figure 6D; Supplementary Figure 5B). Unlike Ramos and Raji cells, this labeling was virtually absent in Daudi cells, consistent with their lower ability to proliferate in the absence of exogenous serine (Figure 6D). In conclusion, these results show that inhibition of PHGDH in the absence of extracellular serine and glycine is able to inhibit purine synthesis and block entry into S-phase of the cell cycle, and promote apoptosis of human lymphoma cell lines.

### Genetic loss and pharmacological inhibition of PHGDH reduces lymphoma progression *in vivo*

To investigate the importance of the SSP pathway in tumor development *in vivo*, we used the Eμ-*MYC* mouse model, which carries a transgene mimicking the t(8:14) translocation of *MYC* and *IGH* characteristic of human Burkitt lymphoma.(45) We first characterized these mice by assessing the expression levels of SSP-related enzymes in B cells isolated from wild-type and lymphoma-bearing *Eμ-Myc* mice. Immunoblotting and IHC showed higher expression of PHGDH and PSAT1 in B cells from lymphoma-bearing *Eμ-Myc* heterozygote mice when compared to wild-type C57BL/6 syngenic mice (Figure 7A-B). When we investigated the ability of *Eμ-Myc* B cells to synthetize serine and glycine *de novo*, we found greater incorporation of labeled U-[^13^C]-glucose in these cells compared to B cells isolated from C57BL/6 mouse spleens (Figure 7C). We then investigated the impact of *Phgdh* deletion on *Myc*-driven tumor development *in vivo*. Deletion of the first enzyme of SSP pathway, *Phgdh*, does not prevent development, but *Phgdh*-deficient mice are born with severe neurological defects and die soon after birth.(46) To overcome these limitations, we first crossed *Eμ-Myc* mice with *Rosa26-CreER^T2/+^mice,(47)* which express a tamoxifen-inducible Cre recombinase enzyme. *Eμ-Myc;Rosa26-CreER^T2/+^* animals were then crossed with *Phgdh^fl/fl^* mice to generate *Eμ-Myc/+;Rosa26-CreER^T2/+^;Phgdh^fl/fl^* mice. Following lymphoma onset, lymphoma cells were purified and allografted by intravenous injection into C57BL/6 syngenic mice. Mice treated in this manner consistently developed splenomegaly and lymphadenopathy from 7 days after injection onwards. The recipients were treated either with vehicle or tamoxifen to excise *Phgdh* specifically in the lymphoma cells (Figure 7D). Notably, knockout of *Phgdh* induced by tamoxifen treatment resulted in a significant reduction in spleen weight in mice sacrificed 20 days after injection (Figure 7E). We next evaluated the pharmacological activity of the PHGDH inhibitor PH-755 on tumor development. C57BL/6 syngenic mice were injected with single transgenic *Eμ-Myc* lymphoma cells and then treated with vehicle or PH-755 (300mg/kg daily, intraperitoneally) for 14 days (Figure 7F). The mice tolerated PH-755 well with no overt side effects from treatment. As with the genetic knockout, pharmacological inhibition of PHGDH with PH-755 also resulted in a significant reduction in lymphoma progression (Figure 7G). Taken together, these data provide evidence that PHGDH is an effective therapeutic target in MYC-driven lymphoma.

**Figure 7.**
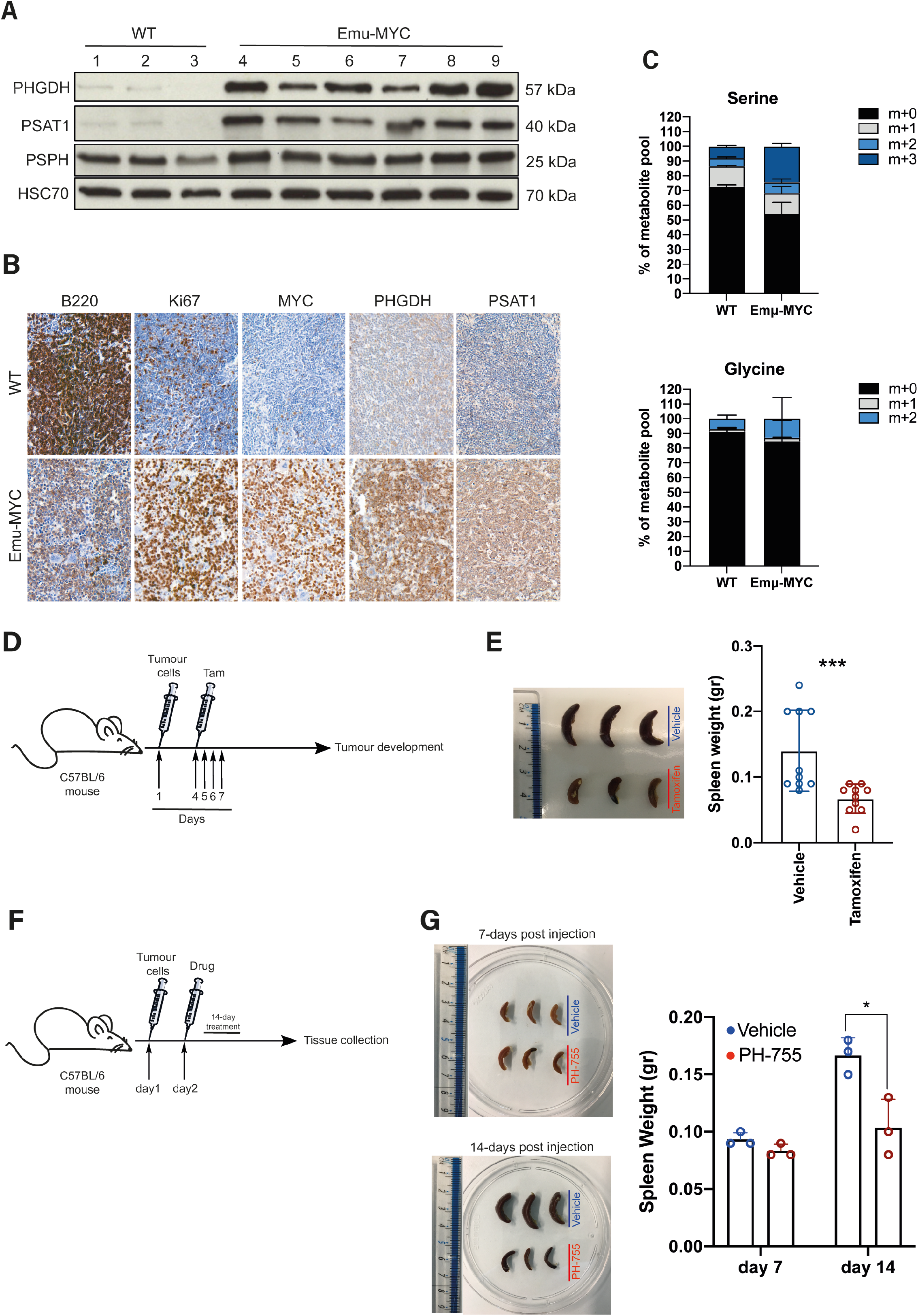
Genetic loss and pharmacological inhibition of PHGDH reduces lymphoma progression *in vivo*. (A) Immunoblot of PHGDH, PSAT1 and PSPH expression in splenic B cells from wild-type (WT; n=3) and *Eμ-Myc* mice (n=6). HSC70 was used as loading control. (B) Representative immunohistochemical staining for B220, Ki67, MYC, PHGDH and PSAT1 abundance in sections of spleens from either wild-type (n=3) or *Eμ-Myc* (n=3) mice. (C) Isotope tracing analysis in splenic B cells isolated from either C57BL/6 WT mice or Eμ-MYC mice and cultured for 2 hours with ^13^C_6_-labeled glucose. Serine, Glycine levels were measured by LC-MS. The percent distribution of each isotopologue of their respective metabolite pool is represented as mean (± SEM) of triplicate cultures and is representative of three independent experiments. (D) Schematic showing allograft lymphoma model, in which lymphoma cells are injected via the tail vein into nine-week-old-male C57BL/6J mice. Three days after lymphoma engraftment, mice are randomized to receive either vehicle or tamoxifen treatment by oral gavage for 4 days. Samples were collected 20 days post-injection. (E) Representative pictures of spleens from mice (n=3 per group) sacrificed 20 days after transplantation (left) and quantification of the spleen weight (right). Data are represented as mean (± SEM); two-tailed Student’s t-test. (F) Schematic showing allograft lymphoma model, in which lymphoma cells are injected via the tail vein into nine-week-old-male C57BL/6J mice. Two days after lymphoma engraftment, mice are randomized to be treated with either vehicle or PH-755 by oral gavage for 14 days. (G) Representative pictures of spleens from mice (n=3 per group) sacrificed after 7 or 14 days post-transplantation (left) and quantification of the spleen weight (right). Data are represented as mean (± SEM); two-tailed Student’s t-test.

## DISCUSSION

Our results establish the importance of the SSP in supporting the proliferation of GC B cells. We show that the SSP is upregulated during B-cell activation and is a metabolic hallmark of the GC reaction. Inhibition of PHGDH, either genetically or pharmacologically, leads to a defect in GC formation and a reduction in high-affinity antibody production. In addition, overexpression of SSP enzymes is a characteristic of GC-derived lymphomas, with high levels of expression predicting a poorer prognosis in DLBCL. In contrast to healthy activated B cells where PHGDH blockade has a cytostatic effect, inhibition of this enzyme in lymphoma cells induces apoptosis, likely reflecting the impact of MYC overexpression. Importantly, inhibition of PHGDH, either specifically within lymphoma cells using an inducible knockout transplantation model, or globally using a pharmacological agent, reduces disease progression highlighting this enzyme as a novel therapeutic target in lymphoma.

These data highlight the interaction between the availability of extracellular serine and the SSP. Collectively these results suggest that the concentration of serine is rate-limiting in GCs, resulting in a requirement for proliferating B cells to synthesize serine from glucose. A recent study has shown serine is an essential metabolite for effector T-cell proliferation.(48) Notably, these authors observed that while effector T cells also upregulate the SSP after activation, they derive the majority of their serine from extracellular sources, with dietary serine availability dictating T-cell responses *in vivo*. Our data suggest an important difference between T- and B-cell biology, due to the anatomical localization of humoral immune responses to the GC. This raises the intriguing possibility that PHGDH inhibition could selectively target the humoral response with relative preservation of T-cell responses, with important implications for the treatment of autoimmune disease and lymphoma.

The role that the SSP plays in cancer is complex and an area of active investigation. Much of the focus has been on the role of PHGDH and its potential as a target for therapy. Notably, while PHGDH does seem to be important for the development for some cancers such as breast cancer, including metastatic disease, observations in other cancer models have suggested that this enzyme is dispensable for tumor development.(23, 27, 49) A previous report suggested that *MYC*-driven lymphomagenesis can occur in the absence of PHGDH.

However, those observations were based on the use of *λ-Myc;Phgdh^+/-^* mice crossed with *Phgdh^+/-^* mice, in a system which generated lymphoma-prone mice with varying expression of PHGDH.(50) It is quite possible that even low levels of PHGDH permit MYC-driven lymphomagenesis, which would explain the differences we observed with our systems. Indeed, *MYC*-driven lymphoma may be particularly susceptible to serine deprivation, as previous work has demonstrated that the progression of lymphoma in Eμ-*MYC* mice can be slowed by reducing the availability of extracellular serine and glycine by dietary restriction.(25) This interaction between extracellular serine and PHGDH inhibition may open up new therapeutic opportunities, either in terms of combining pharmacological inhibition of PHGDH with a serine-glycine free diet, or in treating lymphoma in serine-glycine deplete environments such as the CNS.(36) Tumor cells may become increasingly dependent on *de novo* serine synthesis as the disease progresses, as the increasing mass of cancer cells consumes microenvironmental serine.

Our observations support the hypothesis that it is serine and glycine themselves that are important for lymphoma cell proliferation and survival, as these cells retain the ability to cycle and have low rates of apoptosis when PHGDH was inhibited in serine-glycine replete conditions. Serine appears to be the more important given our observations that culture with glycine and formate only partially rescues the cycling of normal and malignant B cells. In addition, flux through the SSP is also known to play roles in other metabolic process such as redox modulation, α-ketoglutarate production and generation of oncometabolites such as D-2-hydroxyglutarate.(51) Taken together, this work provides important evidence that PHGDH is a viable target for the treatment of pathological B-cell proliferation, either for the modulation of humoral immunity or in lymphoma.

## METHODS

### Animal studies

Mice (3–5 per cage) were allowed access to food and water *ad libitum* and were kept in a 12-hour day/night cycle starting at 7:00 until 19:00. Rooms were kept at 21°C at 55% humidity. Mice were allowed to acclimatize for at least 1 week prior to experimentation. They were then randomly assigned to experimental groups. B6;129P2-PHGDH<tm2Bsi> were purchased from RAKEN BRC (Stock no. RBRC02574). *Eμ-Myc/+* transgenic mice (Stock no: 002728) and *Rosa26-CreER^T2/+^* mice (Stock no: 008463) were purchase from the Jackson Laboratory. *Cd19-Cre KI* mice were kindly provided by Dr. Dinis Calado (The Francis Crick Institute). To test the impact of *Phgdh* deletion on GC response, we generated a mouse model in which *Phgdh* could be conditionally deleted in B cells by crossing *Phgdh^fl/fl^* mice with *Cd19-Cre KI* mice to obtain *Phgdh^fl/fl^;Cd19-Cre* knockout mice. To investigate the impact of *Phgdh* deletion on *Eμ-Myc*-driven tumor development, *Eμ-Myc/+* mice were crossed with the *Rosa26-CreER^T2/+^* mice to generate *Eμ-MYC/+;Rosa26-CreER^T2/+^* mice. *Eμ-Myc/+;Rosa26-CreER^T2/+^* mice were then crossed either with *Phgdh^fl/fl^* or *Phgdh^+/+^* mice to generate *Eμ-Myc/+;Rosa26-CreER^T2/+^;Phgdh^fl/fl^* or *Eμ-Myc/+;Rosa26-CreER^T2/+^;Phgdh^+/+^* genotypes. Following lymphoma onset, lymphoma cells from these mice were purified and used for the adoptive transfer lymphoma experiments.

For T cell-dependent immunization, either *Phgdh^fl/fl^;Cd19-Cre* or *Phgdh^+/+^;Cd19-Cre* mice (7- to 12-week-old) were injected intravenously (i.v) with 1×10^9^ sheep red blood cells (RBC) (TCS BIOSCIENCES LTD, SB054). GCs were analyzed at day 8 after immunization. NP (4-hydroxy-3-nitrophenyl acetyl) was conjugated to CGG (Chicken γ-globulin) at a ratio of NP15-CGG. Mice were injected subcutaneously on the plantar surface of the foot with 20μg NP18-CGG precipitated in alum (ThermoFisher Scientific). For PHGDH inhibitor experiments, C57BL/6J mice (7- to 12-week-old) mice were i.v. injected with 1×10^9^ sheep RBCs. After 24 hours, mice were randomized to receive vehicle (0.5% methylcellulose (Sigma, H7509), 0.5% Tween-80 (Sigma, P8192)) or 300 mgkg^-1^ PH755 (Raze Therapeutics) prepared in vehicle twice a day by oral gavage for the duration of the experiment. GCs were analyzed at day 8 after immunization.

For transplantable lymphoma experiment, *Eμ-Myc* mice harboring lymphomas were euthanized according to licence guidelines. Spleens were collected immediately on ice and homogenized and passed through 70-μm cell strainer in complete RPMI 1640 (Gibco, 21875091) supplemented with 10% fetal calf serum (FCS) (Gibco, 26400044), 2.5% 1M HEPES buffer (Sigma, H0887), 1% GlutaMAX (Gibco, 35050038), 1% Penicillin-Streptomycin (Gibco, 15140122), and 0.1% 0.05M β-mercaptoethanol (Sigma, M3148). Splenocyte suspensions were washed in PBS and incubated in ACK Lysis Buffer (LONZA, 0000879569) for 5 minutes at room temperature (RT) to lyse residual erythrocytes. Lymphoma cells were washed and filtered through 70-μm cell strainer. Cells were resuspended in complete media 2×10^6^ cells in 200μl were injected into C57BL/6J mice (7- to 12-week-old). To investigate the effect of PGHDH inhibition on tumor development, mice were randomized to receive vehicle or 300 mg/kg PH-755 one day after lymphoma injection. Vehicle or drug was administrated once a day by oral gavage for 14 days. Mice were monitored from the conclusion of the treatment to the duration of the experiment. Moribund mice were euthanized according to license guidelines and lymph nodes and spleens were collected for further analysis.

For PHGDH excision experiments, 2×10^6^ *Eμ-Myc/+;Rosa26-CreER^T2/+^;Phgdh^fl/fl^* or *Eμ-Myc/+;Rosa26-CreER^T2/+^;Phgdh^+/+^* cells were injected via tail vein into 7- to 12-week-old C57BL/6J mice. Three days after lymphoma injection, mice were randomized to receive vehicle (sunflower oil; Sigma, S5007) or 3mg at day 1 then 2mg of tamoxifen (Sigma, T5648) daily for 3 days prepared in vehicle. Vehicle or tamoxifen injection was administrated by oral gavage. Mice were monitored every other day for lymphoma development by palpation and euthanized when they reached pre-defined humane endpoints. Lymph nodes and spleens were collected for further analysis.

### Cell culture

#### Cell lines

All the human cell lines used in this study were obtained from Cell Services at the Francis Crick Institute, London, UK. All cell lines underwent routine quality control, which included Mycoplasma detection, STR profiling and species identification for validation. Cells were cultured at 37°C in a humidified atmosphere of 5% CO_2_. DAUDI, RAMOS and NAMALWA cells were cultured in RPMI 1640 medium (Gibco, 21875091) supplemented with 5% FCS; JEKO-1, RAJI, DOHH2 and DG-75 cells were cultured in RPMI 1640 medium (Gibco, 21875091) supplemented with 10% FCS; BJAB and OCI-LY3 cells were culture in RPMI 1640 medium supplemented with 20% FCS; KARPAS 422 and SUDHL4 cells were cultured in ATCC RPMI 1640 medium (Gibco, A1049101) supplemented with 20% and 10% FCS, respectively; GRANTA-519 cells were cultured in DMEM medium (Gibco, 41966052) supplemented with 10% FCS and 2mM L-glutamine; OCI-LY17 cells were cultured in IMDM medium (Gibco, 21980032) supplemented with 20% FCS.

#### Primary cells

Buffy cones were obtained from the UK National Blood Service for investigation of healthy B cells. Human naïve B cells were isolated using the MACSxpress Whole Blood B Cell Isolation Kit (Miltenyi Biotec, 130-098-190) according to the manufacturer’s instructions. Residual erythrocytes were lysed using 1X Red Blood Cell Lysis Buffer (Invitrogen, 00-433). Cells were then washed in full medium consisting of RPMI 1640 (Gibco, 21875091), 10% FCS (Gibco, 26400044), 2.5% 1M HEPES buffer (Sigma, H0887), 1% GlutaMAX (Gibco, 35050038), 1% Penicillin-Streptomycin (Gibco, 15140122), and 0.1% 0.05M β-mercaptoethanol (Sigma, M3148). Cell viability (trypan blue) was >90%. The purity and recovery of the enriched B cells was evaluated by flow cytometry and the proportion of CD19^+^ cells was >95% in all cases.

Resting B cells were stimulated with 20μg/ml Goat F(ab’)2 anti-human IgM + IgG (Stratech Scientific, 109-006-127), 250ng/ml human recombinant MEGACD40L (Enzo, ALX-522-110) and 20ng/ml Recombinant human IL-4 (R&D system, 204-IL-010). Cells were collected at specific time points for analysis.

Murine naïve B cells were purified from spleen by magnetic negative selection by using Pan B Cell Isolation Kit (Miltenyi Biotec, 130-095-813) according to the manufacturer’s instructions and maintained in appropriate culture medium. Typically, purified B cells were plated at a concentration of 1×10^7^/ml and cultured with at 37°C with 5% CO_2_ in RPMI 1640 medium (Gibco, 21875091) supplemented 10% FCS (Gibco, 26400044), 2.5% 1M HEPES buffer (Sigma, H0887), 1% GlutaMAX (Gibco, 35050038), 1% Pencillin-Streptomycin (Gibco, 15140122), and 0.1% 0.05M β-mercaptoethanol (Sigma, M3148). Murine resting B cells were stimulated with 20μg/ml Goat F(ab’)2 anti-mouse IgM + IgG (Stratech Scientific, 115-006-068), 250ng/ml mouse recombinant MEGACD40L (Enzo, ALX-522-120) and 20ng/ml Recombinant mouse IL-4 (R&D system, 404-ML). Cells were collected at specific time points for analysis. For all serine and glycine-deprivation experiments, cells were cultured in standard media deprived of L-serine and L-Glycine.

### Immunoblot analysis

Cells were lysed on ice for 30 minutes using lysis buffer (1% [vol/vol] Nonidet P-40, 20mM Tris-HCl, pH 8.0, 150mM NaCl, and 5mM EDTA with protease inhibitors (Roche diagnostics, Cat. No. 11873580001) and phosphatase inhibitors (sodium fluoride and sodium orthovanadate (Sigma, Cat. No. S7920 and S6508). Samples were centrifuged and the protein content of the supernatant was measured using the Protein Assay Dye Reagent (Biorad, #500-0006). Immunoblotting was performed using from 10 to 50 μg of protein lysate. The following antibodies were used for detecting the following proteins: PHGDH (Cell Signaling Technology, human-specific: #66350; mouse-specific: #13428), PSAT1 (ThermoFisher Scientific, PA5-22124), PSPH (ThermoFisher Scientific, PA5-22003), anti-HSC70 (Santa Cruz Biotechnology, sc-7298). All secondary HRP-conjugated antibodies were a from Cell Signaling Technology. Membranes were exposed using Amersham Hyperfilm ECL (GE Healthcare) and developed using a Curix 60 developer (Agfa). Films were scanned and quantified using ImageJ 1.50c. All values were normalized to the HSC70 loading control, and relative fold-change was calculated with the isotype control antibody treated cells taken as 100% of expression.

### qPCR

Total RNA was isolated from cells using the RNeasy Mini-Kit (QIAGEN, 74104) according to the manufacturer’s instructions. RNA was reverse transcribed using oligo(dT) primers (Promega, C1101), M-MLV (Promega, M1701) in the presence of RNase inhibitor (Promega, N2511). Quantitative PCR was performed on a QuantStudio 12 Flex Real-Time (Applied Biosystems). A standard curve was generated for each human gene from untreated HeLa cells and for each mouse gene from untreated NIH3T3 cells. The average complementary DNA concentration was determined using the standard curve method and made relative to β-Actin. Primers used for qPCR were all purchased from Applied Biosystems and are listed below:

TaqMan probe for human PHGDH: Hs01106329_m1
TaqMan probe for human PSAT1: Hs00795278_mH
TaqMan probe for human PSPH: Hs00190154_m1
TaqMan probe for beta-Actin: Hs01060665_g1
TaqMan probe for human PHGDH: Mm01623589_g1
TaqMan probe for human PSAT1: Mm07293542_m1
TaqMan probe for human PSPH: Mm01197775_m1
TaqMan probe for beta-Actin: Mm02619580_g1

### Immunohistochemistry

All tissues were fixed in 10% neutral buffered formalin and were embedded in paraffin. Tissue microarrays (TMAs) of triplicate 1-mm diameter cores were prepared from paraffin blocks using a manual tissue arrayer (Beecher Scientific) as previously described (52). Cut sections or TMAs were fully drained and placed in a 60°C oven for a minimum of two hours. Slides were then de-paraffinized in xylene and rehydrated using a series of absolute ethanol solutions and distilled water. Heat induced epitope retrieval (HIER) was performed for 10 minutes in a pressure cooker using boiling citric acid-based antigen unmasking solution (Vector, H3300). After retrieval, slides were placed into a wash buffer containing 1X TBS-tween (Agilent Dako, S3306) before immunohistochemical staining. Specific primary antibodies were diluted in Agilent antibody diluent (S080983-2) and incubated for 40 minutes at room temperature. The dilutions and manufacturer details of all the antibodies used are listed in Table 1. The slides were washed and then incubated with biotinylated secondary antibodies (Vector, anti-rat BA-9401, anti-rabbit BA-1000) for 30 minutes at room temperature. The slides were washed again before staining with the VECTASTAIN^®^ Elite ABC peroxidase kit (Vector, PK-6100), in combination with a DAB substrate kit (Vector, SK-4100). The slides were then counterstained with hematoxylin and then scanned with the Pannoramic 250 Flash II system. Immuno-staining was quantified by computerized image analysis using the DensitoQuant tool in Pannoramic Viewer (3DHistTech).

**Table 1.** Details of antibodies used for immunohistochemistry.

| Anti-Mouse antibodies |  |  |  |  |  |  |
| --- | --- | --- | --- | --- | --- | --- |
| Target | Manufacturer | Cat. No. | Dilution | HIER | Protocol | Detection |
| PNA | Vector | B-1075 | 1 in 500 | HIER | Standard | Lectin |
| PHGDH | Thermofisher | PA5-24633 | 1 in 400 | HIER | Standard | Anti-rabbit |
| PSAT1 | Thermofisher | PAS-22124 | 1 in 200 | HIER | Standard | Anti-rabbit |
| B220 | BD | 550286 | 1 in 50 | None | Standard | Anti-Rat |
| MYC | Abcam | ab32072 | 1 in 500 | HIER | Standard | Anti-rabbit |
| Ki67 | Dako | M7249 | 1 in 100 | HIER | Standard | Anti-Rat |
| Anti-human antibodies |  |  |  |  |  |  |
| PHGDH | Sigma Prestige | HPA021241 | 1 in 1000 | HIER | Standard | Anti-rabbit |
| PSAT1 | Thermofisher | PAS-22124 | 1 in 100 | HIER | Standard | Anti-rabbit |

### Flow cytometry

For surface staining, single-cell suspension was washed twice in staining buffer (PBS containing 2% FCS and 2mM EDTA). To detect sheep RBC-induced GC, DZ and LZ B cells, splenocytes were incubated with directly conjugated monoclonal antibodies (Anti-CD19-PE, (BD Bioscience, 7073631), anti-B220-BV510 (BioLegend, 103247), anti-FAS-BV421 (BD Bioscience, 562633), anti-CD38-PE-Cy7 (BioLegend, 102718); anti-CXCR4-Alexa488 (ebioscience, 53-9991-80), anti-CD86-APC (BioLegend, 105012)) and Fixable viability dye eFluor780 staining (ebioscience, 65-0865-14) for 30 minutes on ice protected from light. Cells were then washed and re-suspended in staining buffer and kept at 4°C until analysis. To identify NP-specific plasma cells following NP-CGG immunization, splenocytes were incubated with directly conjugated monoclonal antibodies (Anti-CD19-BUV395, (BD Horizon, 563557), anti-B220-BV510 (BioLegend, 103247), anti-FAS-PerC/P-Cy5.5 (eBioscience, 45-5892-82), anti-CD38-PE-Cy7 (BioLegend, 102718); anti-IgM-APC (ebioscience, 17-5790-82), anti-IgG1-BV421 (BD Horizon, 562580), anti-CD138-BV786 (BD Bioscience, 740880) and anti-NP-PE. To assess PHGDH and PSAT1 levels *in vivo*, splenocytes were first surface stained for GC markers then washed twice in staining buffer and treated with fixation/permeabilization solution Cytofix/Cytoperm (BD Bioscience, 554722) for 30 minutes on ice in the dark. Cells were washed twice in 1X Perm/Wash buffer (BD Bioscience, 554723) and then incubated with either anti-PHGDH (1/250) (Cell Signaling Technology, #13428) or anti-PSAT1 (1/400) (ThermoFisher Scientific, PA5-22124) in staining buffer for 1 hour on ice in the dark and washed twice as previously. Cell were stained with the secondary antibody anti-rabbit AlexaF647 (1/500) (Cell Signaling Technology, #4414) for 30 minutes at room temperature. Cells were then washed, re-suspended in staining buffer and kept at 4°C until analysis. For intracellular Active Caspase-3 staining, cells were stained for surface markers (when required) then fixed/permeabilized by using PE Active Caspase-3 Apoptosis Kit (BD Bioscience, 550914) according to the manufacturer’s protocol. To evaluate cell cycle profile, 10μM BrdU was then added to culture media for 30-45 minutes. Cells were then harvested, fixed and stained with APC anti-BrdU antibody and 7-AAD using the APC BrdU Flow kit (BD Pharmingen, 552598) following the manufacturer’s instructions. Fluorescence was acquired using Fortessa flow cytometer (BD Bioscience) with subsequent analysis using FlowJo software (Tree Star). Analysis was performed after gating on live singlet cells.

### ELISA

Serial dilutions of serum samples were analyzed by ELISA on NP_2_-BSA (5 μg/ml)–coupled microtiter plates to detect high affinity NP-specific IgG1. Alkaline phosphatase (AP)-conjugated primary antibodies anti-IgG1 (Southern Biotech) were developed with p-nitrophenyl phosphate dissolved in Tris buffer (SIGMAFAST, Sigma-Aldrich). The absorbance was measured at 405 nm, plotted against dilution and relative antibody titres were read as the dilution where absorbance reached an arbitrary threshold.

### Metabolite extraction and LC-MS

Cells were seeded at 5×10^6^ per well serine/glycine-free complete media for 1 hour. After 1h, cells were transferred to serine/glycine free media containing ^13^C-U-glucose (2g/L) (Cambridge Isotope Laboratories, CLM-1396-PK) for additional 2 hours. Cells were then washed with PBS before being metabolically quenched by transferring to dry ice. For PHGDH inhibition experiment, cells were pretreated with 10μM PH-755 (Raze Therapeutics) diluted in DMSO or DMSO alone for 1 hour before labelling with ^13^C-U-glucose. Metabolites were extracted by resuspending the cell pellet in ice-cold HPLC-grade methanol, acetonitrile and H2O at volume ratio 50:30:20 for 1 hour at 4°C with three sonication steps (8 minutes each step) within the hour. After sonication, samples were centrifuged for 10 minutes at 14,000rpm and the supernatant was collected. The extraction solvent was dried in a glass insert placed in a LC-MS vial (Agilent 5182-0716) using a SpeedVac (Christ RVC 2-33 CDplus.). Then, 20 μl of metabolite extraction buffer containing 5μM ^13^C, ^15^N-Valine (used as an internal extraction standard) were added to each dried sample and metabolite samples were stored at −80°C for subsequent analysis. Triplicates of identically seeded and treated cells were analyzed.

Metabolite analysis was performed by Liquid chromatography–mass spectrometry (LC-MS) using a Q-Exactive Plus (Orbitrap) mass spectrometer (Thermo Fisher Scientific) coupled with a Vanquish UHPLC system (Thermo Fisher Scientific). The chromatographic separation was performed on a SeQuant^®^ Zic^®^pHILIC (Merck Millipore) column (5μm particle size, polymeric, 150 x 4.6 mm) using a gradient program at a constant flow rate of 300 μl/min over a total run time of 25 min. The elution gradient was programmed as decreasing percentage of B from 80% to 5% during 17 minutes, holding at 5% of B during 3 minutes and finally re-equilibrating the column at 80 % of B during 4 minutes. Solvent A was 20 mM ammonium carbonate solution in water supplemented by 4 ml/L of a solution of ammonium hydroxide at 35% in water and solvent B was acetonitrile. MS was performed with positive/negative polarity switching using a Q Exactive Orbitrap (Thermo Scientific) with a HESI II probe. MS parameters were as follows: spray voltage 3.5 and 3.2 kV for positive and negative modes, respectively; probe temperature 320°C; sheath and auxiliary gases were 30 and 5 arbitrary units, respectively; and full scan range: 70–1,050 m/z with settings of AGC target and resolution as balanced and high (3 × 10^6^ and 70,000), respectively. Data were recorded using Xcalibur 4.2.47 software (Thermo Scientific). Mass calibration was performed for both ESI polarities before analysis using the standard Thermo Scientific Calmix solution. To enhance calibration stability, lock-mass correction was also applied to each analytical run using ubiquitous low-mass contaminants. Parallel reaction monitoring (PRM) acquisition parameters were the following: resolution 17,500; collision energies were set individually in HCD (high-energy collisional dissociation) mode. Metabolites were identified and quantified by accurate mass and retention time and by comparison to the retention times, mass spectra, and responses of known amounts of authentic standards using TraceFinder 4.1 EFS software (Thermo Fisher Scientific). Label incorporation and abundance was estimated using TraceFinder 4.1 EFS software. The level of labelling of individual metabolites was estimated as the percentage of the metabolite pool containing one or more ^13^C atoms after correction for natural abundance isotopes. Abundance was given relative to the internal standard.

### Single-cell RNA sequencing analysis of human tonsillar B-cell subsets

Processed single-cell RNA sequencing gene expression datasets from tonsillar immune cells were obtained (37) and analyzed in Seurat (v3) (53) using previously determined cell type annotations. Imputation of gene expression counts for visualisation was performed with MAGIC (54).

### Statistics

Statistical significance was assessed Prism 9.1.1 software (GraphPad Software, Inc.). Groups were compared using an unpaired t test, Mann Witney U test or ANOVA with Tukey’s post hoc test depending on the number of groups and distribution. For survival comparison a logrank test was used. Graphs show the mean and standard error of the mean (SEM). A *P* value of less than or equal to 0.05 was considered to indicate statistical significance. When analyzing variables with more than 2 categories, P values were adjusted for multiple comparisons. \**p* < 0.05, \*\**p* < 0.01, \*\*\**p* < 0.001, \*\*\*\**p* < 0.0001 in the figures.

### Study approval

All animal studies were conducted in compliance with UK Home Office approved licences (Animals (Scientific Procedures) Act 1986 and the EU Directive 2010). Animal experiments were subject to ethical review by the Francis Crick Animal Welfare and Ethical Review Body and carried out under UK Home Office project licence P319AE968. Lymphoma samples were obtained from Barts Cancer Institute tissue bank, London, UK. Ethical approval was confirmed by the East London & The City Health Authority Local Research Ethics Committee (Reference number 10/H0704/65), and written informed consent was also obtained in accordance with the Declaration of Helsinki.

## AUTHOR CONTRIBUTIONS

AD designed and performed the experiments, analyzed and interpreted data, and wrote the manuscript; MT and ECC designed experiments, analyzed and interpreted data; NL and JM designed and performed the metabolomic experiments, analyzed and interpreted the LC-MS data; HWK and LJ analyzed and interpreted the single-cell RNA sequencing datasets; PC analyzed and interpreted the statistical data, AG and LZ designed the flow cytometry and ELISA experiments, analyzed and interpreted data; JGG and DPC designed experiments and edited the manuscript, KHV designed experiments, analyzed and interpreted the data, edited the manuscript, and supervised the study, and JCR designed and performed the experiments, analyzed and interpreted the data, wrote and edited the manuscript, and supervised the study. All authors approved the final submission.

## ACKNOWLEDGEMENTS

The authors are very grateful to all of the patients who generously donated their tissue to the tissue biobank. We also thank Kamil Kranc for his thorough review of the manuscript. This work was supported by fellowships and grants from the Wellcome Trust (110020/Z/15/Z to JCR; 213555/Z/18/Z to HWK), Cancer Research UK (C596/A26855 to KHV). In addition, this work was supported by The Francis Crick Institute which receives its core funding from Cancer Research UK (FC001557, FC001057), the United Kingdom Medical Research Council (FC001557, FC001057), and the Wellcome Trust (FC001557, FC001057), and by Barts Cancer Institute which receives funding from a Cancer Research UK Centre Grant (C355/A25137). For the purpose of Open Access, the authors have applied a CC BY public copyright licence to any Author Accepted Manuscript version arising from this submission.

## Notes

**CONFLICTS OF INTEREST** KHV is on the board of directors and a shareholder of Bristol Myers Squibb, a shareholder of GRAIL, Inc., and on the science advisory board (with stock options) of PMV Pharma, RAZE Therapeutics and Volastra Therapeutics, Inc. She is also on the SAB of Ludwig Cancer. KHV is a co-founder and consultant of Faeth Therapeutics. She has been in receipt of research funding from Astex Pharmaceuticals and AstraZeneca and contributed to CRUK Cancer Research Technology filing of Patent Application WO/2017/144877. JGG has received research funding from Celgene, Janssen and Acerta and honoraria from Abbvie, Acerta, Celgene, Gilead, Janssen, Novartis, Pharmacyclics and Roche. The other authors have declared that no conflict of interest exists.

### Competing Interest Statement

KHV is on the board of directors and a shareholder of Bristol Myers Squibb, a shareholder of GRAIL, Inc., and on the science advisory board (with stock options) of PMV Pharma, RAZE Therapeutics and Volastra Therapeutics, Inc. She is also on the SAB of Ludwig Cancer. KHV is a co-founder and consultant of Faeth Therapeutics. She has been in receipt of research funding from Astex Pharmaceuticals and AstraZeneca and contributed to CRUK Cancer Research Technology filing of Patent Application WO/2017/144877. JGG has received research funding from Celgene, Janssen and Acerta and honoraria from Abbvie, Acerta, Celgene, Gilead, Janssen, Novartis, Pharmacyclics and Roche. The other authors have declared that no conflict of interest exists.

