## Supplementary data for "PHGDH is required for germinal center formation and is a therapeutic target in *MYC*-driven lymphoma"

**SUPPLEMENTARY FIGURES**

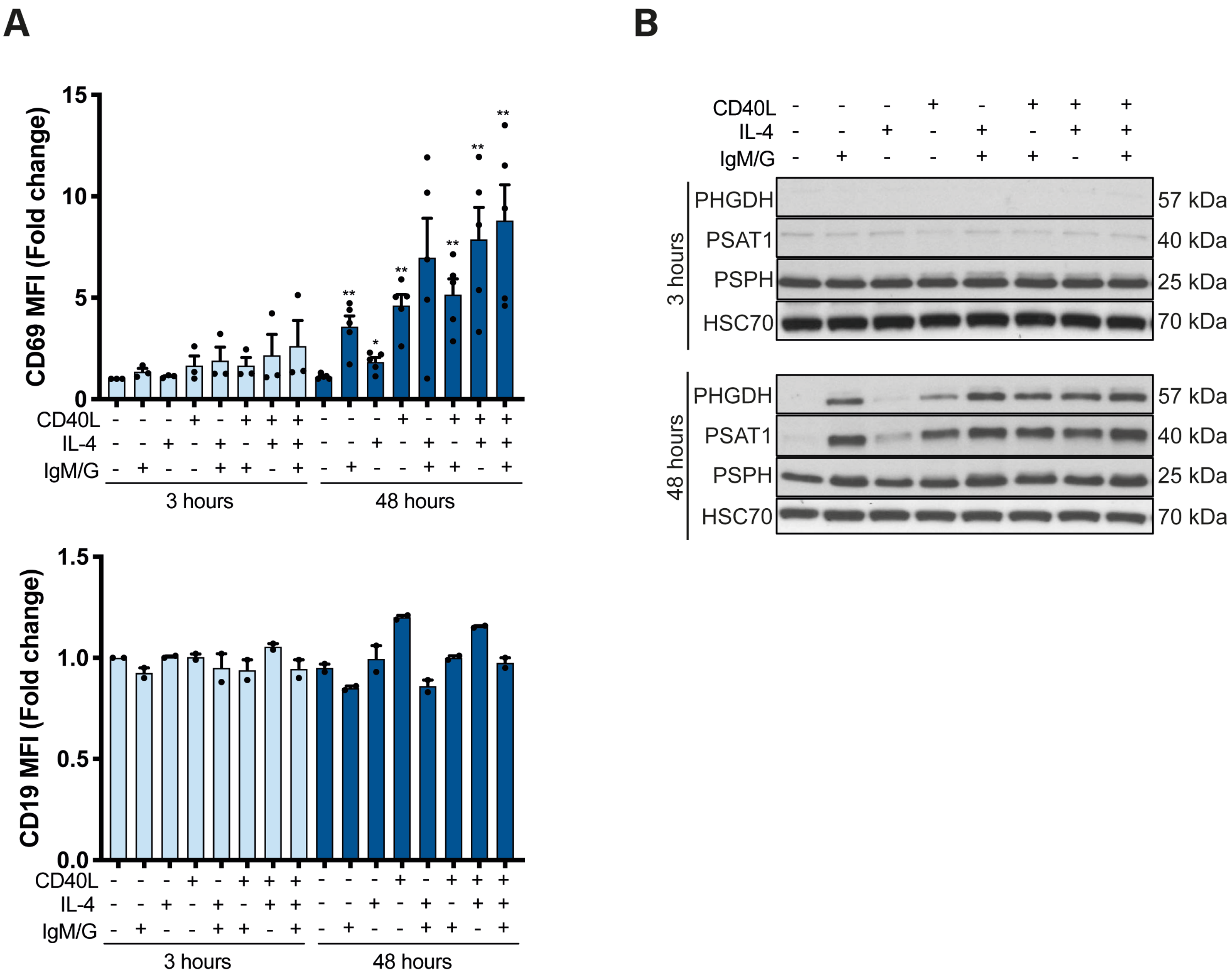


**Supplementary Figure 1. Activation of human B cells leads to upregulation of the serine synthesis pathway.**

(A) Flow cytometric analysis of surface expression of CD69 (above) and CD19 (below) and (B) representative immunoblots of PHGDH, PSAT1 and PSPH proteins levels in resting human mature naïve B cells (-) or in B cells stimulated with anti-IgM/G, CD40L and or IL-4 for 3 and 48 hours (+). Individual samples (dots) and means (bars) values are plotted (**p* < 0.05, ***p* < 0.01, ****p* < 0.001, *****p* < 0.0001; Mann-Whitney test).

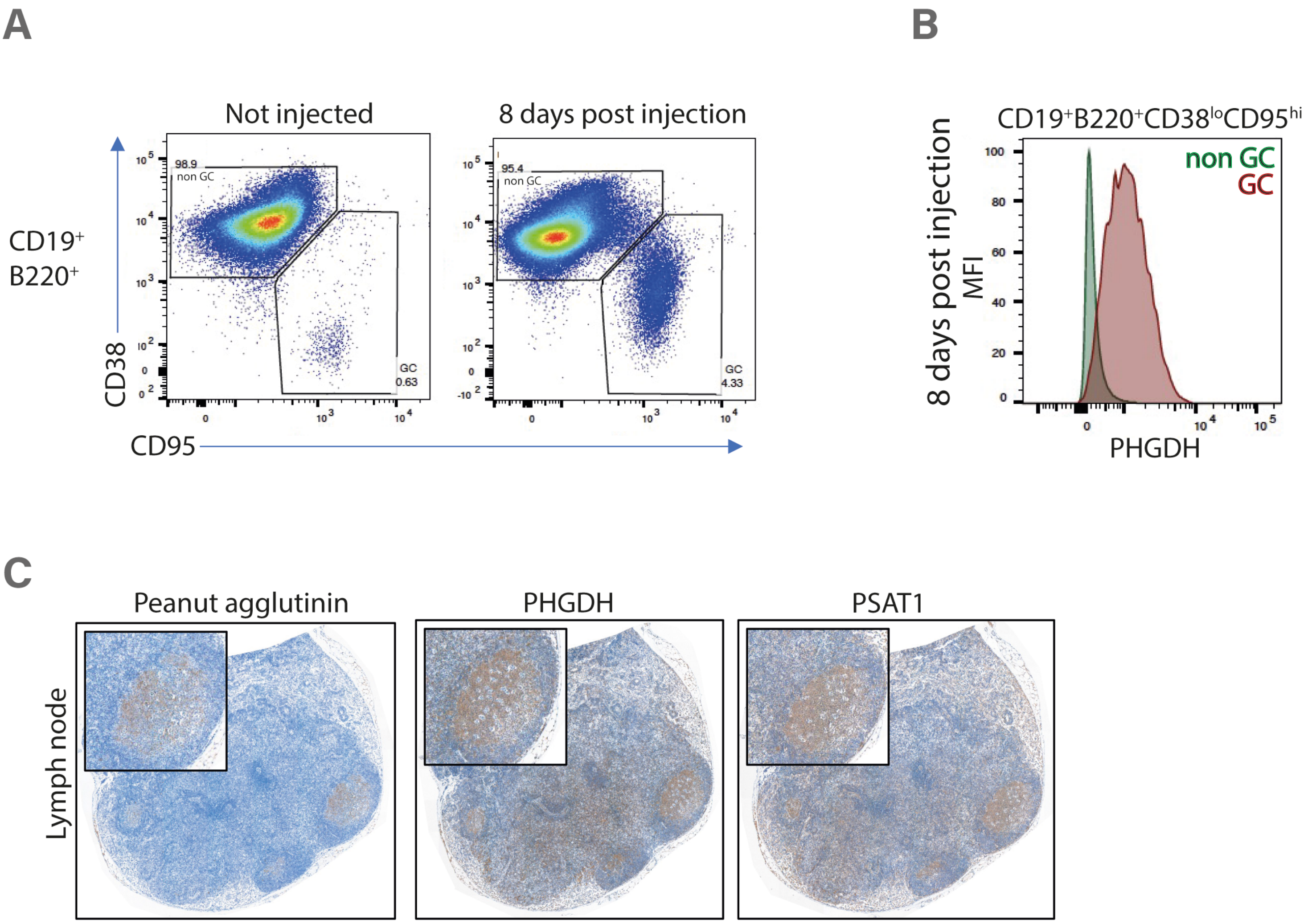


**Supplementary Figure 2. In vivo activation of serine synthesis pathway in GC B cells**

(A) Representative flow cytometric analysis of splenic B cells in BL6 mice left unimmunized (left panel) or immunized with sheep RBCs for 8 days (right panel) to identify GC B cells ($CD19^{+}B220^{+}CD38^{lo}CD95^{hi}$). (B) Representative flow cytometric analysis of PGHDH expression within GC B cells ($CD19^{+}B220^{+}CD38^{lo}CD95^{hi}$) isolated from BL6 mice immunized with sheep RBCs for 8 days. (C) Representative immunohistochemical staining for peanut agglutinin (PNA) as GC marker, PHGDH and PSAT1 on consecutive sections derived from mice lymph nodes 8 days after sheep RBC immunization. Data are representative of three independent experiments.

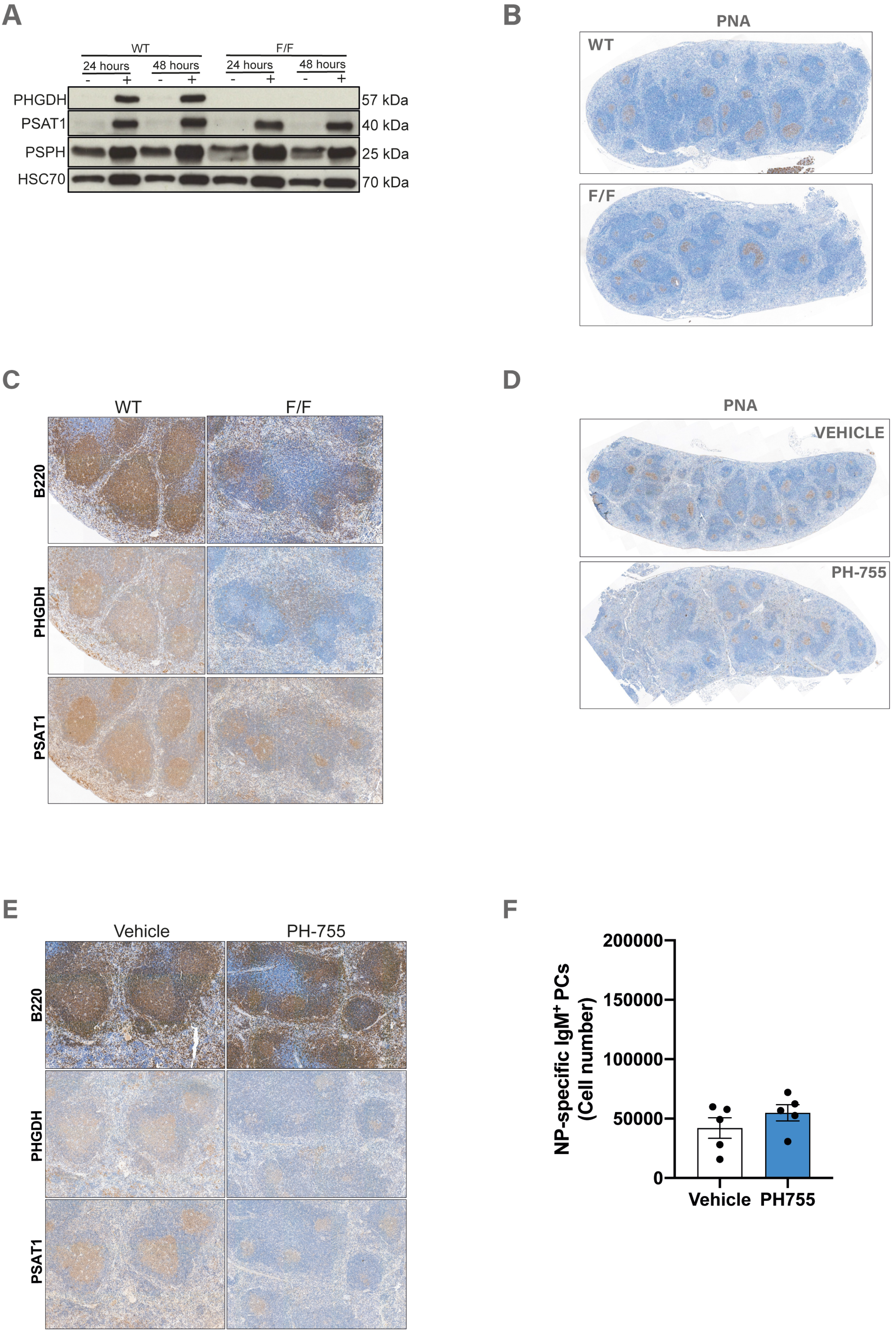
A
WT
F/F
24 hours
48 hours
24 hours
48 hours
- + - + - + - +
PHGDH
PSAT1
PSPH
HSC70
57 kDa
40 kDa
25 kDa
70 kDa
B
PNA
WT
F/F
C
WT
F/F
B220
PHGDH
PSAT1
D
PNA
VEHICLE
PH-755
E
Vehicle
PH-755
B220
PHGDH
PSAT1
F
NP-specific IgM+ PCs
(Cell number)
200000
150000
100000
50000
0
Vehicle
PH755

**Supplementary Figure 3. Inhibition of PHGDH impairs GC responses**

(A) Analysis of PHGDH, PSAT1 and PSPH protein levels in B220$^+$ B cells isolated from either *PHGDH$^{+/+}$/CD19-Cre* (WT) or *PHGDH$^{fl/fl}$/CD19-Cre* (F/F) immunized with sheep BC for 8 days, and treated in vitro with (+) or without (-) anti-IgM antibody, CD40 ligand (CD40L) and interleukin-4 (IL-4) for 24 and 48 hours before protein extraction. (B) Representative immunohistochemical staining for PNA in spleen sections derived from *Phgdh$^{+/+}$;Cd19-Cre* (WT) and *Phgdh$^{fl/fl}$;Cd19-Cre* (F/F) mice 8 days after immunization with sheep RBCs. Data are representative of three independent experiments. (C) Representative immunohistochemical staining for B220, PHGDH and PSAT1 on consecutive spleen sections derived from mice described in (B). (D) Representative immunohistochemical staining for PNA in spleen sections from mice injected with sheep RBC and treated with PH-755 or vehicle control. Data are representative of three independent experiments. (E) Representative immunohistochemical staining for B220, PHGDH and PSAT1 on consecutive spleen sections derived from mice described in (D). Data are representative of three independent experiments. (F) Numbers of NP-specific IgM$^+$ PCs (total number per popliteal lymph nodes). Wild-type BL6 mice were treated with either Vehicle (n=5) or PH-755 (n=5) for 7 days. Animals were injected with NP-CGG one day before the beginning of PH-755 treatment. Popliteal lymph nodes were collected 8 days post NP-CGG immunization.

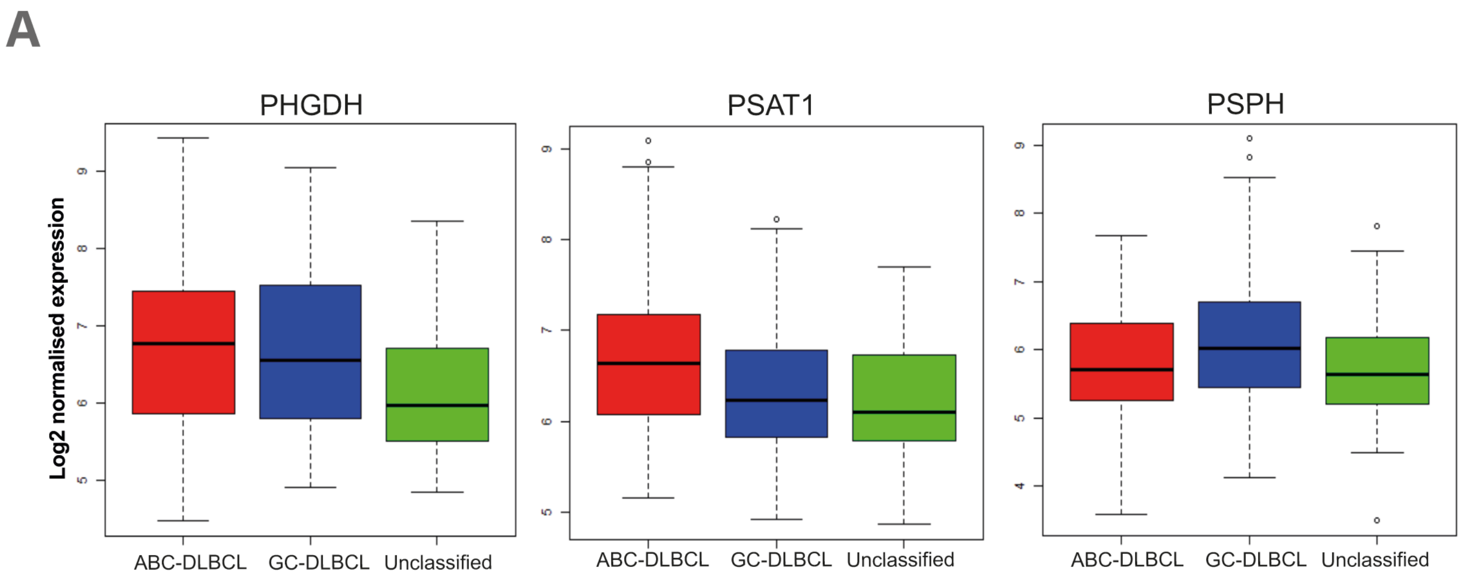


**Supplementary Figure 4. No difference in expression of SSP enzymes in DLBCL subtypes**

(A) The expression of PHGDH, PSAT1, PSPH mRNA was analyzed in a published dataset (GSE10846) to compare the expression of these enzymes in activated B-cell (ABC) like, germinal centre B-cell (GCB) like, or unclassified DLBCL.

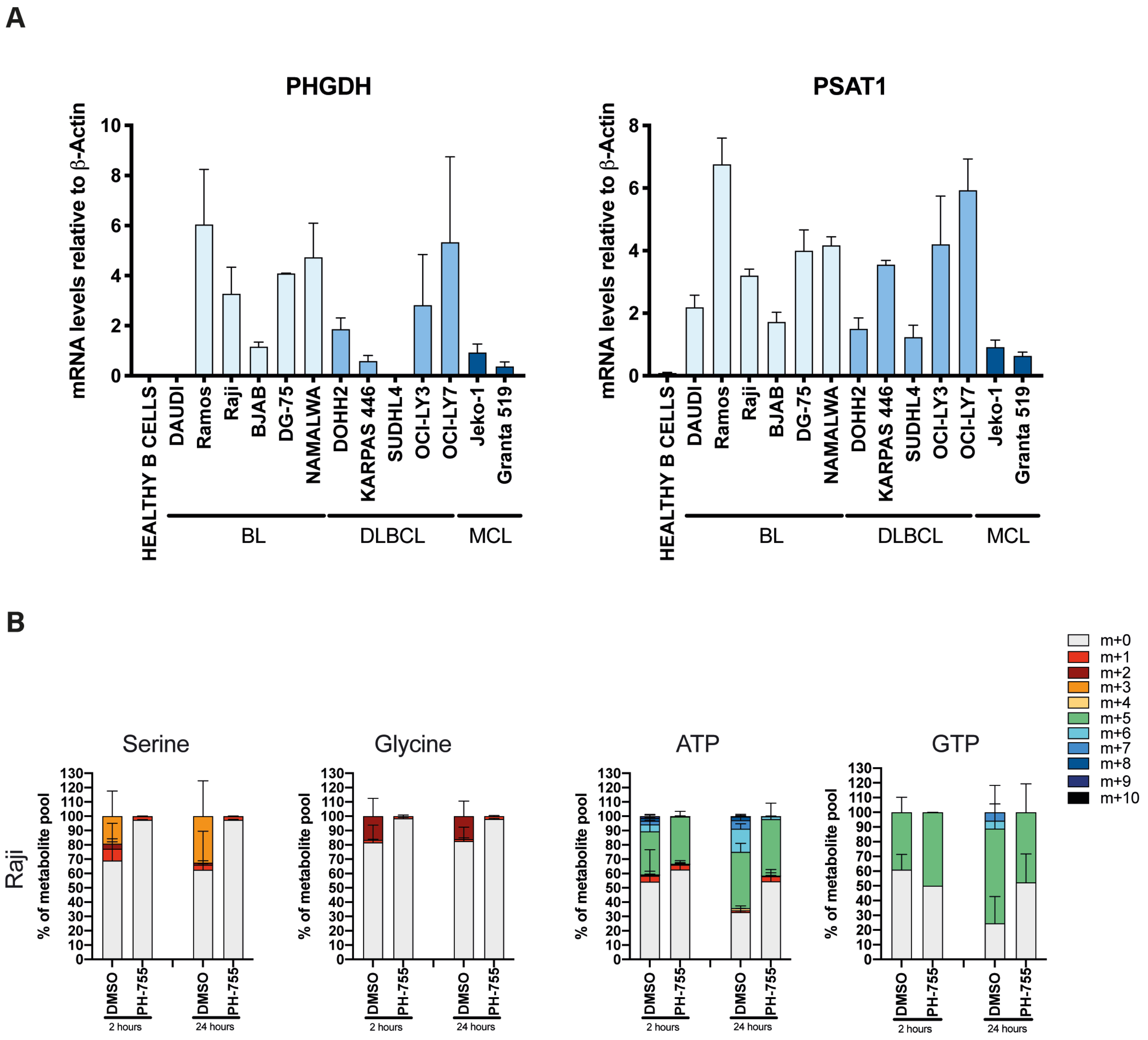


**Supplementary Figure 5. Serine synthesis pathway is activated in in Burkitt lymphoma cells**

(A) Relative mRNA expression of SSP enzyme genes from different human B-cell lymphoma lines compared with primary B cells. Cell line were grouped into three groups (BL, DLBCL and mantle cell lymphoma (MCL) cell lines). Data are presented as mean (± SEM) of two independent experiments. (B) Mass isotopologue distribution of U-[$^{13}C_6$]-glucose-derived serine, glycine, ATP and GTP for Raji cells cultured for 2 and 24 hours in medium lacking serine and glycine in presence of U-[$^{13}C_6$]-glucose (10mM) and treated with DMSO or 10µM PH-755. Serine, Glycine, ATP and GTP levels were measured by LC-MS. The percentage distribution of each isotopologue for their respective metabolite pool is shown. Data are presented as mean (± SEM) of six repeats and are representative of three independent experiments.
